# NMR unveils an N-terminal interaction interface on acetylated-α-synuclein monomers for recruitment to fibrils

**DOI:** 10.1101/2020.08.17.254508

**Authors:** Xue Yang, Baifan Wang, Cody L. Hoop, Jonathan K. Williams, Jean Baum

**Affiliations:** Department of Chemistry and Chemical Biology, Rutgers University, Piscataway, New Jersey 08854

**Keywords:** Amyloid formation, Amyloid seeding, IDP–IDR interactions, Aggregation inhibition, Parkinson’s Disease

## Abstract

Amyloid fibril formation of α-synuclein (αS) is associated with multiple neurodegenerative diseases, including Parkinson’s Disease (PD). Growing evidence suggests that progression of PD is linked to cell-to-cell propagation of αS fibrils, which leads to seeding of endogenous intrinsically disordered monomer via templated elongation and secondary nucleation. A molecular understanding of the seeding mechanism and driving interactions is crucial to inhibit progression of amyloid formation. Here, using relaxation-based solution NMR experiments designed to probe large complexes, we probe weak interactions of intrinsically disordered acetylated-αS (Ac-αS) monomers with seeding-competent Ac-αS fibrils and seeding-incompetent off-pathway oligomers to identify Ac-αS monomer residues at the binding interface. Under conditions that favor fibril elongation, we determine that the first 11 N-terminal residues on the monomer form a common binding site for both fibrils and off-pathway oligomers. Additionally, the presence of off-pathway oligomers within a fibril seeding environment suppresses seeded amyloid formation, as observed through thioflavin-T fluorescence experiments. This highlights that off-pathway αS oligomers can act as an auto-inhibitor against αS fibril elongation. Based on these data taken together with previous results, we propose a model in which Ac-αS monomer recruitment to the fibril is driven by interactions between the intrinsically disordered monomer N-terminus and the intrinsically disordered flanking regions (IDR) on the fibril surface. We suggest that this monomer recruitment may play a role in the elongation of amyloid fibrils and highlight the potential of the IDRs of the fibril as important therapeutic targets against seeded amyloid formation.

**Significance:** Cell-to-cell spreading of αS fibrils leads to amyloid seeding of endogenous monomer. Detailed atomic-level mechanistic understanding of the fibril seeding process of αS is essential for design of therapeutic approaches against Parkinson’s disease. In light of its complexity, this process remains ill-defined at the molecular level. Using relaxation-based solution NMR experiments, we mapped a common N-terminal binding interface of the Ac-αS intrinsically disordered monomer with Ac-αS fibrils and off-pathway oligomers to elucidate critical monomer–aggregate interactions during seeded aggregation and in equilibrium with mature aggregates. From this work, we propose a new paradigm, in which Ac-αS monomer recruitment to the fibril is driven by interactions between the intrinsically disordered monomer N-terminus and the flanking IDRs on the fibril surface.

## 1. Introduction

Amyloid formation of the intrinsically disordered protein (IDP) α-Synuclein (αS) is closely associated with the pathogenesis of a variety of neurodegenerative disorders, including Parkinson's disease (PD), dementia with Lewy bodies, and multiple-systems atrophy (1). The pathology of αS is still not clearly understood, as multiple species along the fibril aggregation pathway, including on-pathway oligomeric intermediates and end-stage fibrils, have been demonstrated to be toxic to neurons (2–8). Abrogation of these toxic species is extremely challenging, in part due to the amyloid seeding process, which is a complex mechanism by which endogenous monomers interact with existing fibril seeds to facilitate additional fibril formation. The mechanism involves multiple microscopic steps, including elongation of fibril ends and surface-mediated secondary nucleation (9–11). Furthermore, disease progression has become increasingly linked to prion-like cell-to-cell propagation of αS aggregates, in which αS oligomers and fibrils are transmitted to neighboring neurons (12, 13). Here, the invading aggregates seed amyloid formation of endogenous monomers to accelerate production of toxic species (3). Thus, disruption of the process by which αS aggregates seed proliferative amyloid formation has become an intriguing target for inhibition of pathological αS self-assembly and may reveal new therapeutic approaches against PD.

Under amyloid fibril formation conditions in vitro, αS monomers self-assemble along multiple pathways to mature into various species of stable off-pathway oligomers, transient on-pathway oligomers that convert into amyloid fibrils, fragmented fibril seeds and mature fibrils, creating a complex, heterogeneous environment (Figure 1a). While the inherent structural inhomogeneity and transient nature of on-pathway oligomers makes them extremely challenging to isolate and characterize, off-pathway oligomers are stable and can be isolated by size exclusion chromatography (SEC). Multiple species of stable, off-pathway αS oligomers under various conditions have been previously isolated (5, 14–16). Lorenzen et al. isolated two forms of stable αS oligomers from amyloid promoting conditions and determined the structure of the smaller oligomers to consist of ~30 monomers oriented in an ellipsoid shape, with an outer cloud of flexible and disordered molecules, based on small angle X-ray scattering (SAXS) (14) (Figure 1a). How these stable oligomers impact amyloid seeding and growth is not well understood. Determining how stable oligomers influence amyloid aggregation mechanisms may provide insight into pathological seeding processes and aid in discovery of novel therapeutic approaches against amyloid seeding.

**Figure 1.**
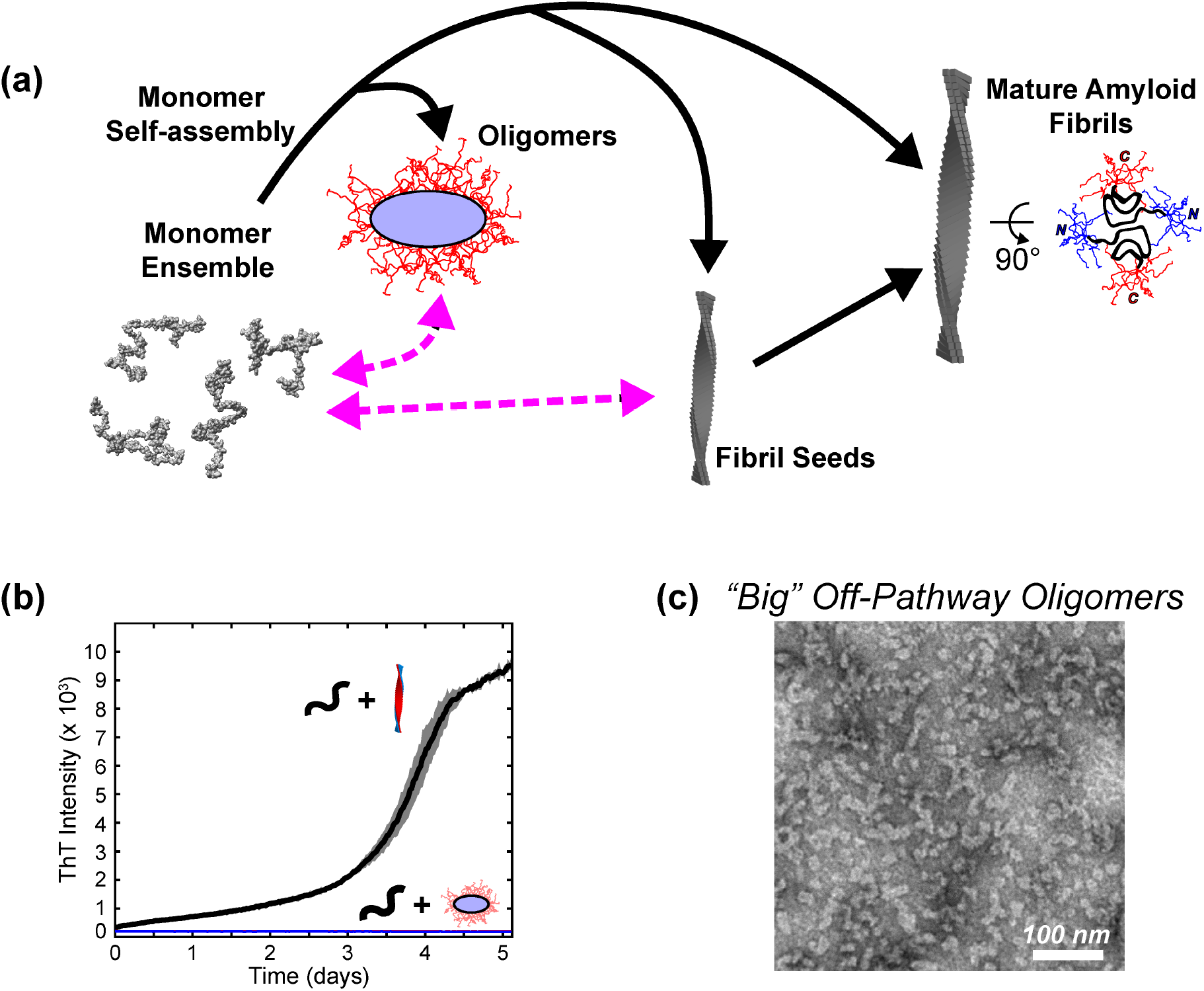
In fibril forming conditions, αS undergoes multiple aggregation pathways. (a) Schematic of the multiple assembly pathways and products formed by αS. In vitro, αS monomers can self-assemble in the same conditions into multiple species of oligomers or into amyloid fibrils, creating a complex heterogeneous environment. Stable oligomers may be formed off-pathway and do not proceed to amyloid formation. Fibril formation is accelerated through fibril seeding processes, such as templated elongation and secondary nucleation. Here, we investigate αS monomer interactions with off-pathway pre-formed oligomers (PFOs) or pre-formed fibrils (PFFs) (magenta, dashed arrows) and the impact of co-existing off-pathway PFOs on kinetics of fibril seeding. The monomer ensemble is based on PED00024 (73). Stable oligomers formed in fibril formation conditions have been shown to have a disordered C-terminal cloud (~55 residues, red) that surrounds the core consisting of the N-terminal and NAC regions (14). Mature amyloid fibrils also have a surface of disordered N-(blue) and C-termini (red) that flank a rigid core (model adapted from PDB: 6h6b (20)). (b) In an in vitro ThT fibril seeding assay at 37°C, Ac-αS PFFs induce the formation of new amyloid fibrils (black- 40 μM monomer, 1 μM fibril seeds). However, big PFOs do not have the ability to induce formation of new amyloid fibrils (blue- 40 μM monomer, 1 μM big PFOs). (c) A TEM image of isolated big PFOs displays their worm-like morphology.

It has been observed previously that the rigid amyloid core can act as a template to seed fibril formation (17, 18), yet the role of intrinsically disordered flanking regions in fibril seeding is not well understood. Structural models derived from solid-state NMR and cryo-electron microscopy (EM) have revealed that αS fibrils consist of two protofilaments with a rigid “Greek key” core flanked by intrinsically disordered N- and C-termini (~40 residues each) (19–23) (Figure 1a). Current inhibition methods primarily target the rigid core. These include using chaperones to block fibril ends or fibril surfaces (9), treating with natural products that promote fibril clustering to reduce fibril fragmentation and hide binding sites for monomer addition (24, 25), and creating high affinity interactions with amyloidogenic monomers to sequester monomers from self-assembly (26). Recently, it has become increasingly more important to understand the role of the intrinsically disordered regions (IDRs) in fibril growth in order to determine their potential for amyloid inhibition. These IDRs are a substantial fraction of the fibril structure and make up a “fuzzy coat” that surrounds the rigid core (19–21). The flexible flanking regions in several amyloid proteins have been found to be highly involved in biological interactions, including binding to receptors, chaperones, membranes, or RNA (27). In αS, modifying the “fuzzy coat” through pathologically relevant N- and C-terminal truncations was shown to modulate the aggregation and seeding propensity of αS fibrils and influence the resulting fibril morphologies (28–30). This suggests that the terminal IDRs are also actively involved in the αS aggregation mechanism. In order to more fully understand the seeding process, it is critical to identify the monomer–fibril interactions, and whether they involve the rigid core or the flanking IDRs.

The weak, transient nature of the monomer–fibril interactions that occur during fibril seeding make it a challenge to identify the key residues involved. Solution NMR is unique in its ability to detect perturbations to soluble proteins at atomic-level resolution. Sophisticated solution NMR experiments, dark state exchange saturation transfer (DEST), have been designed to probe the exchange of NMR-visible, free monomers with the surface of large, NMR-invisible complexes, such as amyloid fibrils, in residue-specific detail not accessible by other techniques. These powerful NMR methods were introduced by probing the interactions between NMR-visible amyloid-β (Aβ) monomers with large, NMR-invisible Aβ protofibrils implicated in amyloid formation (31). More recently, studies have used DEST to determine site specific interactions of IDPs with membranes (32), lipopolysaccharides (33), unilamellar vesicles (34), and in self-assembly (35, 36). Our lab has recently used ^15^N-DEST to determine the interaction interfaces on immunoglobulin protein β_2_-microglobulin (β_2_m) for collagen I, which facilitates β_2_m amyloid formation (37).

Here we use NMR to characterize monomer–fibril and monomer–oligomer interactions that occur under conditions promoting templated elongation in order to gain a molecular understanding of the interaction between the N-terminally acetylated-αS (Ac-αS) monomer and these complex aggregates. Through relaxation-based NMR experiments, we find that Ac-αS pre-formed fibrils (PFFs) and off-pathway pre-formed oligomers (PFOs) are in a dynamic equilibrium with NMR-visible monomers. ^15^N-DEST experiments and ^15^N-transverse relaxation measurements reveal that the N-terminal 11 residues of Ac-αS monomers interact with PFFs and stable off-pathway PFOs, suggesting that the binding interface of the monomer is similar for both fibrils and oligomers. When added to a monomer solution prepared under templated elongation conditions, these same PFOs significantly delay fibril formation in a concentration dependent manner and can act as auto-inhibitors of amyloid formation when co-existing with monomers and PFFs. Our results suggest that delays in fibril formation by PFOs may be explained by competing monomer N-terminal interactions between PFFs and PFOs through their common binding modes. Identification of this shared N-terminal binding site on Ac-αS monomers for fibrils and off-pathway oligomers, taken together with previous data, leads us to propose that the N-terminus of the intrinsically disordered monomer interacts with the intrinsically disordered terminal flanking regions (IDR) of the fibrils and oligomers, a shared feature of these aggregate structures. These IDP–IDR interactions may play a critical role in recruiting monomer to the fibril under conditions that promote templated elongation. While the structured fibril core is known to be critical to the templated assembly of amyloid fibrils, it is important to consider the role of the N- and C-terminal fibril IDRs in fibril seeding and assembly and their potential as targets against amyloid propagation.

## 2. Results

### Stable Ac-αS oligomers formed under amyloid promoting conditions do not convert to fibrils or seed fibril growth

We investigate the interactions of αS monomers with seeding-competent αS pre-formed fibrils (PFFs) or seeding-incompetent αS off-pathway pre-formed oligomers (PFOs) using NMR experiments. All αS used in these studies is N-terminally acetylated, as the N-terminus of αS is post-translationally acetylated in vivo (2). N-terminal acetylation increases the helical propensity of the N-terminus, enhances membrane binding (38, 39), and has been shown to influence fibril formation kinetics and morphologies of fibrils produced (40). Thus, interactions of N-terminally acetylated-αS (Ac-αS) monomers with seeding-competent PFFs and seeding-incompetent PFOs is physiologically significant to understand seeding mechanisms.

We isolated two species of stable Ac-αS oligomers by SEC (Figure S1a), which were prepared under fibril formation conditions, at 37°C with 300 rpm shaking for 5 h. Notably, these stable Ac-αS oligomers do not convert to fibrils and do not seed amyloid formation of Ac-αS monomers in conditions that favor templated elongation (PBS, pH 7.4, 37°C) within 5 days (Figure 1b). Thus, while they are formed in conditions that promote fibril formation, these oligomers are formed separately from the fibril formation pathway, and so are defined as “off-pathway.” In isolation they do not progress to amyloid fibrils upon incubation at 37°C for 5 days, as evidenced by the lack of thioflavin T (ThT) fluorescence enhancement (Figure S1c) and maintain their morphologies without evidence of fibril formation even after 42 days as assessed by TEM (Figure S1f–g). In addition, these stable oligomers are not converted to fibrils by incubation with pre-formed Ac-αS fibril seeds in templated elongation conditions over five days, as detected by ThT (Figure S1d). Both species of stable Ac-αS oligomers show a similar distribution of secondary structures by circular dichroism, with reduced β-sheet content relative to mature fibrils (Figure S1b). However, they are distinct from each other in size and morphology. “Big” oligomers show an overall elongated, worm-like morphology as visualized by transmission electron microscopy (TEM) (Figure 1c), while “small” oligomers are uniformly circular in shape (Figure S1e). Both big and small oligomers are similar in size and morphology to non-acetylated αS oligomer species previously isolated and characterized by the Otzen group (14). Taken together, these results establish these isolated big and small pre-formed oligomers (PFOs) as off-pathway species that can be used as negative controls of seeding.

### Off-pathway Ac-αS oligomers are non-toxic to SH-SY5Y cells

Extensive work has been done to investigate the observed toxicity of αS fibrils, and while an exact understanding of the toxicity mechanism has not been reached, αS amyloid fibrils are thought to exert their toxic effect through interactions with multiple different cellular pathways (19, 41–43). Oligomers have also shown significant levels of cellular toxicity (8, 44, 45). MTS (3-(4,5-dimethylthiazol-2-yl)-5-(3-carboxymethoxyphenyl)-2-(4-sulfophenyl)-2H-tetrazolium) cell viability assays were used to evaluate the potential toxicity of the big and small PFOs used in this work. After incubating SH-SY5Y neuronal cells with Ac-αS PFFs for 24 h, cell viability is reduced in a dose dependent manner (Figure 2a, gray). However, when treating SH-SY5Y cells with the either big or small PFOs (up to 5 μM monomer equivalent) for 24 h, no significant MTS reduction is observed (Figure 2a, red, blue). To ensure an accurate assessment of fibril or oligomer entry into the cells, SH-SY5Y cells were treated with PFFs or PFOs covalently linked with ATTO-488. After 24 h incubation, cells were stained with DAPI and purified mouse anti-αS antibody (Figure 2b–c, S2). Cells treated with either PFFs or PFOs showed cellular entry of the aggregates (Figure 2b–c, S2). PFF treated cells show 3.3 fold higher anti-αS fluorescence relative to monomer control cells (Figure 2b–c, S2), indicating that fibrils or other aggregates efficiently accumulate in the cellular environment. However, cells treated with big or small PFOs only show 1.34 fold and 1.76 fold higher anti-αS fluorescence relative to monomer treated cells (Figure 2b–c, S2), respectively. Thus, these off-pathway oligomers do not induce significant accumulation of αS aggregates in the cellular environment.

**Figure 2.**
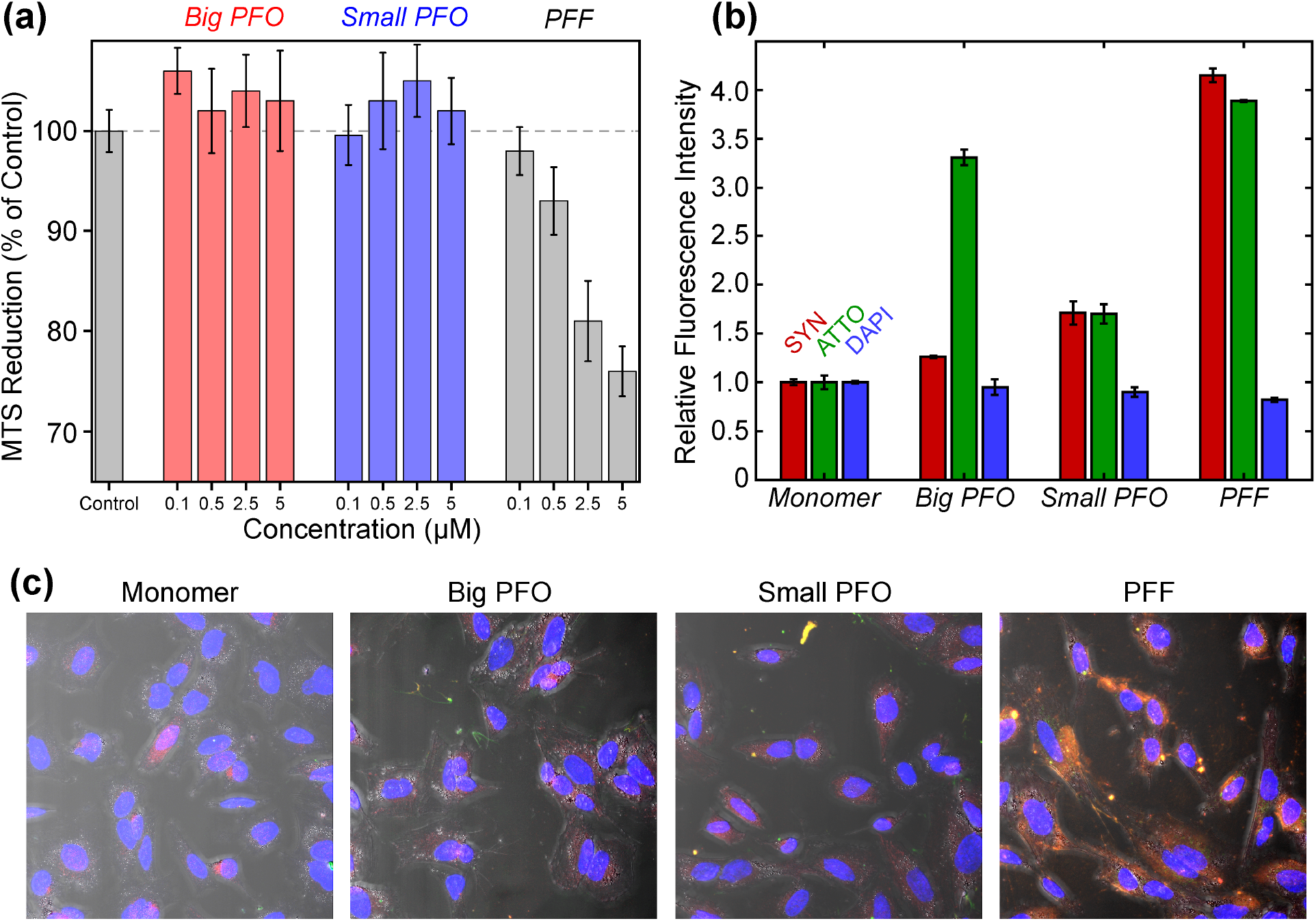
Off pathway oligomers are not toxic to SH-SY5Y cells. (a) Cell viability assessed using an MTS reduction assay. Ac-αS monomer was used as the control. % MTS reduction for all aggregate species was normalized to the level of the control (dashed line). Both big (red) and small (blue) PFOs show no reductions in cell viability with increasing concentrations relative to monomers. Conversely, increasing concentrations of Ac-αS PFFs show a dose-dependent reduction in cell viability. (b) Quantitation of the relative fluorescence intensities observed from the (c) confocal fluorescence images of SH-SY5Y cells treated with monomer, PFF, big PFO, or small PFO and then stained with anti-αS-antibody (red), ATTO-488 dye (green), or DAPI (blue). In (c) the composite images are shown, which are an overlay of the three anti-αS-antibody, ATTO-488, and DAPI-treated channels along with the DIC image showing the outline of the cell body. Separate confocal images of anti-αS-antibody, ATTO-488, DAPI channels and the DIC images are shown for monomer, PFF, big PFO, and small PFO treated cells in Figure S2.

### Mature Ac-αS fibrils and off-pathway oligomers are in a dynamic equilibrium with NMR-visible monomers

Ac-αS PFFs and PFOs were characterized by solution NMR in order to gain a molecular understanding of an equilibrium between Ac-αS monomers and fibrils/oligomers. In seeding experiments, fibrils are sonicated to reduce the heterogeneity of fibril lengths (Figure S3a, d–f) and increase the number of fibril ends to promote elongation (Figure S3b) (46). A ^1^H–^15^N HSQC spectrum of sonicated PFFs (TEM shown in Figure S4a) shows indistinguishable chemical shifts from Ac-αS monomers under the same conditions (Figure 3a, S5d). Because the fibril species are large and tumble slowly in solution, they are not detected in solution NMR. However the presence of observable monomer signals indicates a detectable population of dissociated monomers that are in equilibrium with the fibrils.

**Figure 3.**
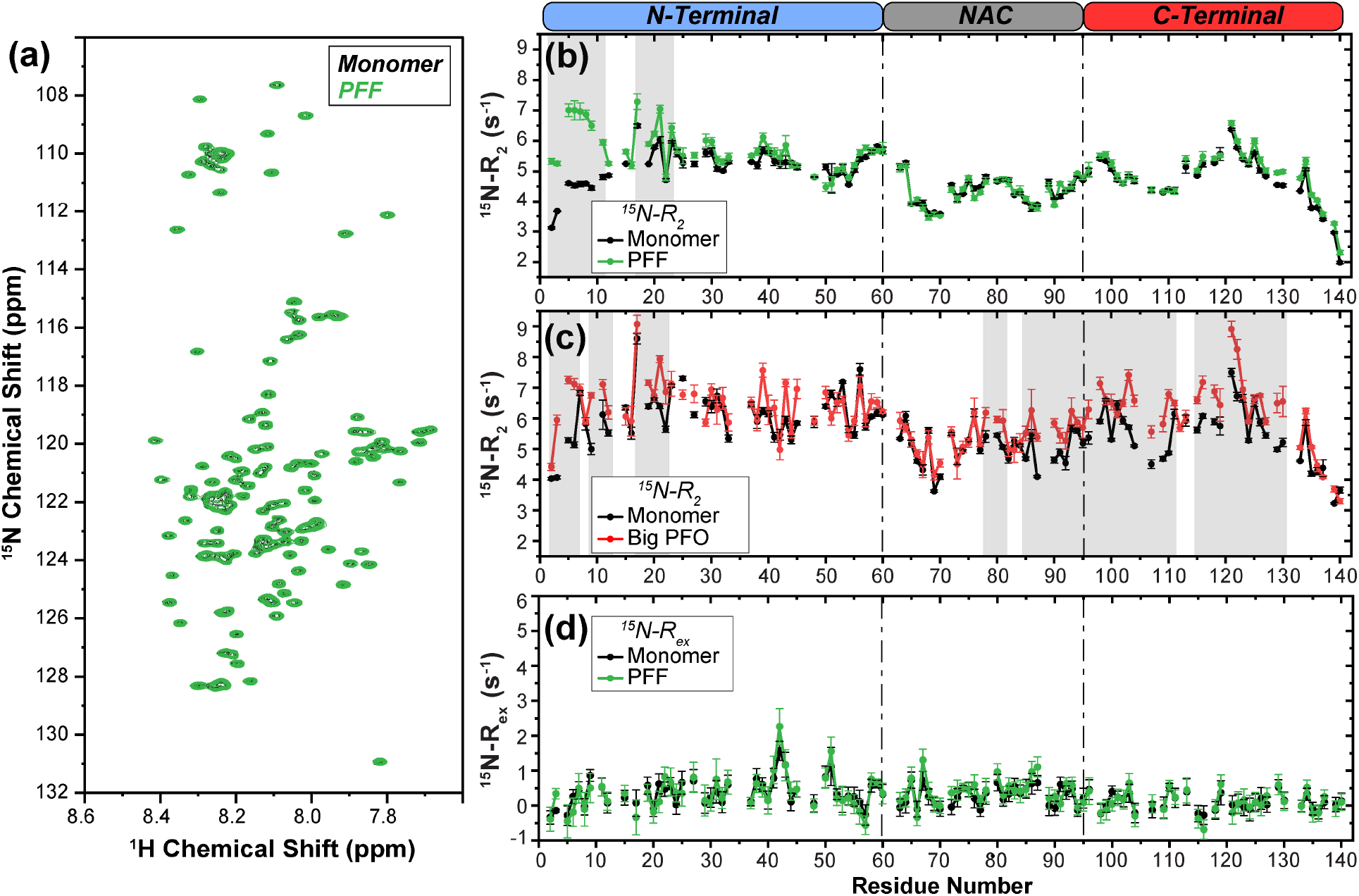
Sonicated fibrils and off-pathway oligomers are in equilibrium with monomers. (a) 1H–^15^N HSQC spectra of 350 μM [U-^15^N]-Ac-αS monomer (black) and 350 μM [U-^15^N]-Ac-αS PFFs (green). An overlay of these spectra shows indistinguishable chemical shifts and linewidths, indicating that the detected species in the [U-^15^N]-PFFs sample is actually monomer dissociated from fibrils, referred to as equilibrium monomer. (b,c) Residue-level ^15^N-transverse relaxation rates (^15^N-R_2_) of (b) 350 μM [U-^15^N]-Ac-αS monomers (black) and 350 μM [U-^15^N]-Ac-αS PFFs (green) and (c) 350 μM [U-^15^N]-Ac-αS monomers (black) and 350 μM [U-^15^N]-Ac-αS big PFOs (red). The three sequential domains of αS are indicated on top (N-terminal, blue; NAC, gray; and C-terminal, red). Dashed lines separate the domains within the plots. Regions of three or more consecutive residues that have a ΔR_2_ between the monomer and aggregate samples ≥ 0.5 s^−1^ are highlighted in gray. (d) Relaxation exchange rates (^15^N-R_ex_) as measured by ^15^N-R_2_ Hahn echo experiments of 350 μM [U-^15^N]-Ac-αS monomers (black) and 350 μM [U-^15^N]-Ac-αS PFFs (green). No substantial contribution of R_ex_ is observed in either sample. The error bars in the relaxation data are propagated from the single exponential decay fitting errors of the underlying ^15^N-R_2_ data. All NMR data were acquired in 10 mM PBS, pH 7.4 with 10% D_2_O and 4°C at 800 MHz ^1^H Larmor frequency.

Similarly, SEC and NMR experiments indicate that both big and small PFOs in solution are also in equilibrium with monomers. After being isolated by SEC, big PFOs were immediately re-injected onto the SEC column, and a monomer peak re-emerged (Figure S4b). This indicates that oligomers are continuously in equilibrium with monomers, and monomers cannot be completely separated from the system. These PFOs also show chemical shifts indicative of the intrinsically disordered Ac-αS monomers in their ^1^H–^15^N HSQC spectra (Figure S5e–f). However, normalization of the peak intensities of the off-pathway oligomer relative to the monomer at the same concentration reveal that big and small PFOs have increased intensity in C-terminal residues (Figure S6a). This is in agreement with solution NMR studies of αS oligomers by the Otzen group, which indicated that residues 86–140 remain mobile, based on comparisons of peak intensities between monomeric and oligomeric samples (47). The increased intensity observed in the C-terminus of our big and small PFOs may be due to superposition of peak intensity resultant from both monomeric and oligomeric species, suggesting that the C-termini make up the “fuzzy” surface of the oligomer, as seen in previous models (Figure 1a) (14, 16).

### A common N-terminal interaction hotspot on Ac-αS monomers is shared between off-pathway oligomers and fibrils

Having established that PFFs and PFOs are in equilibrium with NMR-visible Ac-αS monomers in solution, we next determined which specific residues of the equilibrium monomers are participating in the monomer–PFF or monomer–PFO interactions using ^15^N NMR relaxation experiments. No chemical shift perturbation of equilibrium monomer from PFFs or PFOs relative to isolated free monomer in solution is detected, indicating weak interactions (Figure 3a, S5d–f). Equilibrium monomers from PFFs show substantially elevated ^15^N-transverse relaxation rates (R_2_) in the N-terminal region (first 23 residues), especially in the early N-terminus (residues 2–11) (Figure 3b, S6b). An increase in ^15^N-R_2_ is indicative of reduced backbone motions on the picosecond–nanosecond timescale due to site-specific direct interactions and/or slowing of global molecular tumbling, or to enhanced conformational exchange on the microsecond–millisecond timescale. The contribution of conformational exchange was assessed by ^15^N in-phase Hahn echo experiments (Figure 3d), which estimate the exchange contribution to the observed ^15^N-R_2_ as ^15^N-R_ex_. No significant R_ex_ is observed in either the free monomer alone or equilibrium monomer from PFFs, suggesting that the enhanced ^15^N-R_2_ values arise from site-specific interactions.

Surprisingly, the same N-terminal region (first 22 residues) was found to have increased ^15^N-R_2_ in equilibrium monomers from PFOs relative to free monomers, with additional increased ^15^N-R_2_ in the late NAC (residues 79–82 and 86–95) and C-terminus (residues 96–111 and 115–130) (Figure 3c, S6b). Taken together with the increased peak intensity in the C-terminus and the Otzen model of a disordered C-terminus in non-acetylated αS oligomers (47), these data are consistent with a superposition of oligomer and monomer C-terminal signals in solution. Because of this, the contribution to the overall peak intensities and apparent ^15^N-R_2_ from these two species is difficult to deconvolute. The intrinsically disordered segments of the oligomer would exhibit slower molecular tumbling as part of the oligomeric complex relative to monomer free in solution. This would result in a globally increased ^15^N-R_2_ of the intrinsically disordered segments of the oligomer relative to that of the monomer. However, the N-terminus of the oligomer is expected to be rigid, buried within the oligomeric core (14, 47) and thus its contribution to peak intensities or ^15^N-R_2_ is likely undetectable by solution NMR. Therefore the elevated ^15^N-R_2_ in the N-terminus of the PFO samples instead indicates perturbation of equilibrium monomer N-terminal backbone dynamics. This suggests that equilibrium monomers from PFOs interact with the oligomers at the same N-terminal binding site as identified for the monomer–PFF interaction at equilibrium.

### Visualizing the Ac-αS monomer–fibril interaction interface using NMR ^15^N DEST experiments

To probe the specific molecular interactions of the NMR-visible equilibrium monomer with NMR-invisible high molecular weight species (fibrils) we use ^15^N-dark state exchange saturation transfer (DEST) (31, 48) experiments. This technique takes advantage of the broad, undetectable resonances of high molecular weight “dark state” complexes (e.g. fibrils) to selectively saturate these species at frequencies far off-resonance and transfer the saturation to an observable “light state” (e.g. unbound equilibrium Ac-αS monomer) in exchange with the complex (31, 48). These experiments are sensitive to slow exchange processes on the ~10 ms– ~1 s timescale (48). A key indicator of this exchange regime is a detectable difference in ^15^N-R_2_ values between a sample with only the free monomer and one containing the dark state. Based on considerations of sensitivity and maximum ΔR_2_ values (~2.5 s^−1^, Figure S6b), the Ac-αS monomer–PFF equilibrium system is the most amenable to ^15^N-DEST experiments. The superposition of equilibrium monomer and oligomer signals and the lower maximum ΔR_2_ (~2.0 s^−1^, Figure S6c–d) in the monomer–PFO equilibrium system makes analysis of the ^15^N-DEST experiments more complicated.

We performed ^15^N-DEST experiments on free ^15^N-Ac-αS monomers in the absence of fibrils (i.e. absence of a dark state) and ^15^N-Ac-αS PFFs in equilibrium with monomers. Spectra of NMR-visible equilibrium monomers (light state) show residue-specific peak intensity loss at multiple distant ^15^N-offsets relative to free monomers in the absence of a dark state. This is indicative of residues in exchange with the dark fibril-bound state and results in broadening of the DEST saturation profile for those residues (Figure 4a–c). This broadening effect is expressed as ΔΘ in Figure 4a. We find that the per-residue DEST saturation profiles of the equilibrium monomer from PFFs is broadened for early N-terminal (first 12) residues (Figure 4a–c). This indicates that indeed the equilibrium monomer is associating with the fibril, forming the detected dark-state complex, and that this association is strongest in the first 12 N-terminal residues.

**Figure 4.**
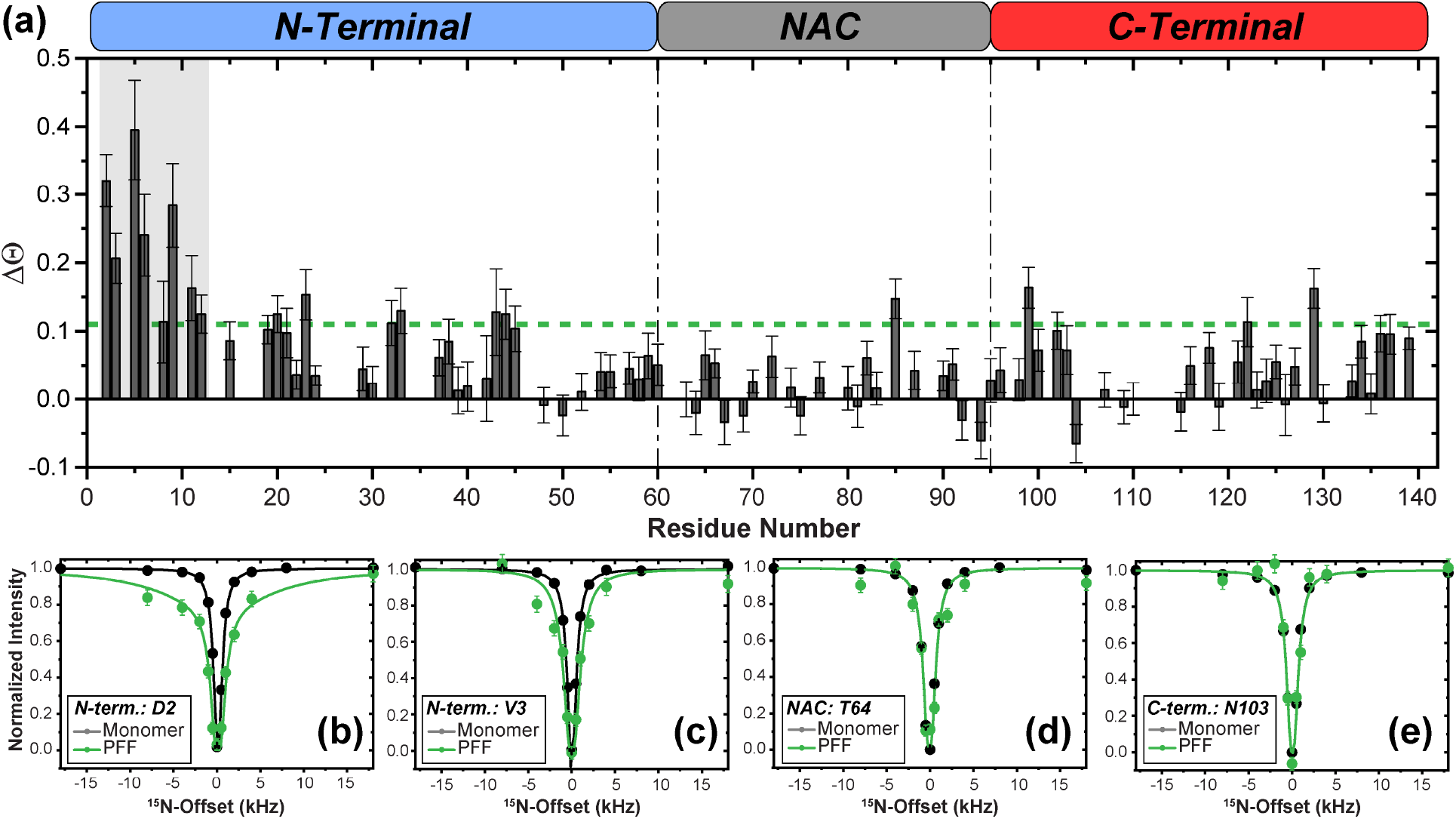
Equilibrium monomers interact via their N-terminus with sonicated fibrils. (a)ΔΘ calculated as the difference in Θ between [U-^15^N]-Ac-αS PFFs and [U-^15^N]-Ac-αS monomers derived from ^15^N-DEST intensities at ±30 kHz and ±1 kHz ^15^N-offsets with a 350 Hz saturation frequency. Regions of consecutive residues with ΔΘ higher than 0.11 (green dashed line), where broadening of the residue-specific DEST profiles is observed, are highlighted in gray. The three linear αS domains are indicated on the top. 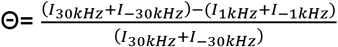; ΔΘ = Θ_+fibrils_ − Θ_−fibrils_. (b–e) Examples of ^15^N-DEST profiles with 350 Hz saturation frequency of 350 μM [U-^15^N]-Ac-αS monomer (black) or 350 μM [U-^15^N]-Ac-αS PFFs (green). Experimental data points are shown as symbols. Fit curves using a two-state exchange model are shown as solid lines. N-terminal residues D2 and V3 show broadening of the DEST profiles in the [U-^15^N]-PFFs sample (presence of a dark state) (b–c), whereas residues in the NAC (T64) and C-terminal (N103) do not show this effect (d–e). Error bars are derived from the noise of the underlying 2D spectra. All experiments were conducted in 10 mM PBS, pH 7.4 with 10% D_2_O, 4°C at 800 MHz ^1^H Larmor frequency.

Representative DEST saturation (350 Hz saturation) profiles for residues in each of the three domains of free Ac-αS monomer in the absence of a dark state or equilibrium Ac-αS monomer are provided in Figures 4b-e. For residues D2 and V3 in the N-terminus, a clear broadening of the profile is observed for the equilibrium monomer relative to free monomer (Figure 4b–c). However, for residues T64 in the NAC and N103 in the C-terminus, minimal differences are observed between the two samples (Figures 4d–e). A simple two-state model that accounts for one light state (unbound equilibrium monomer) and one dark state (equilibrium monomer bound to fibril) fits to residue-specific ΔR_2_ values (R_2_^equilibrium^-R_2_^free monomer^) and DEST profiles with 350 Hz and 150 Hz RF saturation (Figure S7, Figure 4b–e). Based on this two-state model, 99.79 ± 0.05% of the monomer is unbound while the remainder is bound to the PFFs with an apparent first order rate constant (k_on_^app^) of 2.36 ± 0.08 s^−1^. Despite this low population of PFF-bound monomer, substantially higher R_2_^bound^ values on the order of 10^3^–10^5^ s^−1^ (Figure S7b) are estimated for the first 11 residues relative to the remainder of the sequence. The analysis of the ^15^N-DEST saturation profiles, calculation of ΔΘ, and simulated estimations of R_2_^bound^ values suggests that within the dynamic monomer–PFF equilibrium there exists a small population of monomers that are in direct contact with the PFFs via a N-terminal hotspot that includes the first 11 residues. N-terminal residues that exhibit increased ^15^N-R_2_ values but do not show the DEST effect are likely not in direct contact with the PFFs, but may still be impacted by the interaction due to their proximity to the binding site. Taken together, the differential ^15^N-R_2_ data and ^15^N-DEST on ^15^N-Ac-αS PFFs suggest that they are in dynamic equilibrium with dissociating and re-associating Ac-αS monomers and that the monomer–fibril direct contacts involve the early monomer N-terminus, which is potentially also an interaction domain for off-pathway oligomers.

### In a kinetic seeding environment, Ac-αS monomers interact with off-pathway oligomers and fibrils at the same N-terminal interaction hotspot

In order to identify monomer residues that interact with pre-formed fibril seeds in a kinetic seeding environment, in which the concentration of monomeric Ac-αS is much higher than those of the PFFs or PFOs, and to compare them with the monomer–PFF interaction interface observed in equilibrium, we probed for residue-specific interactions of ^15^N-Ac-αS monomers in the presence of low concentrations of PFFs or PFOs under conditions that promote templated elongation (PBS at pH 7.4). No chemical shift perturbations are observed in the ^1^H–^15^N HSQC spectra of Ac-αS monomers upon addition of any of the aggregates (Figure S5a–c). Interestingly, upon addition of either PFFs or PFOs, we observe perturbation of the monomer backbone dynamics, with the largest increases in ^15^N-R_2_ observed in the N-terminal nine residues (Figure 5a,b), consistent with the equilibrium results above. However, the maximum ΔR_2_ observed is ~ 1 s^−1^, indicating a slower exchange process than what is observed in the equilibrium systems. These slow processes preclude accurate quantitation by DEST measurements. Since the ^15^N-R_2_ perturbation occurs in the same N-terminal region of the monomer as in the equilibrium systems without conformational exchange contribution (Figure 5c), we expect that the increased ^15^N-R_2_ values result from monomer interactions specifically at the N-terminal hotspot with the PFFs or PFOs in a seeding environment that promotes templated elongation.

**Figure 5.**
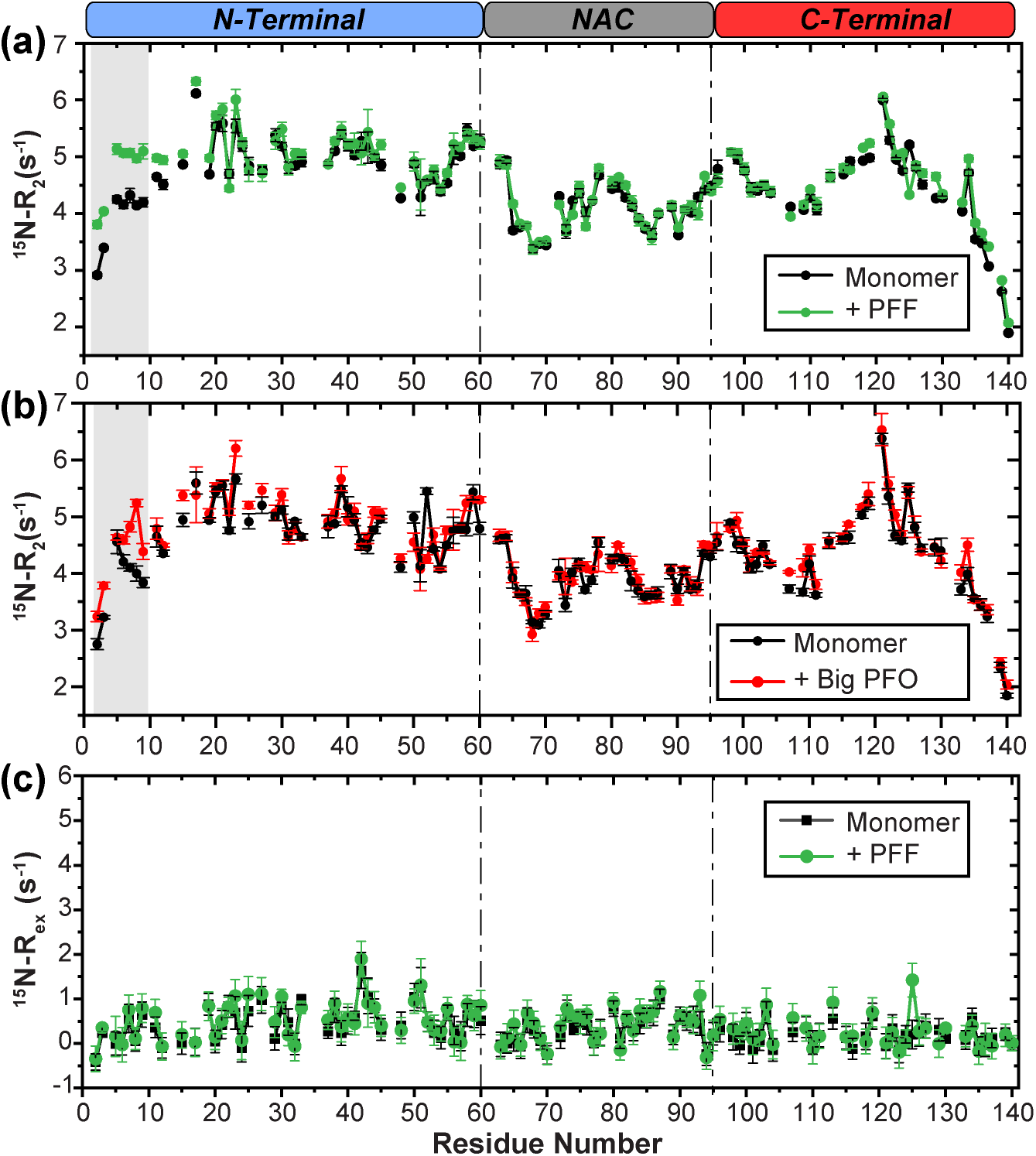
Perturbation to [U-^15^N]-Ac-αS monomers in the presence of PFFs or PFOs in a kinetic seeding environment reveals a common interaction interface. Residue-level ^15^N-R_2_ of (a) green: 90 μM [U-^15^N]-Ac-αS monomer + unlabeled PFFs (1:1 molar ratio) and black: 90 μM [U-^15^N]-Ac-αS monomer alone, and (b) red: 90 μM [U-^15^N]-Ac-αS monomer + unlabeled big PFOs (1:1 molar ratio) and black: 90 μM [U-^15^N]-Ac-αS monomer alone. Regions of three or more consecutive residues that have a ΔR_2_ between monomers in the presence and absence of aggregates ≥ 0.5 s^−1^ are highlighted in gray. (c) ^15^N-R_ex_ as measured by ^15^N-R_2_ Hahn echo experiments of 90 μM [U-^15^N]-Ac-αS monomers in the absence (black) or presence of 90 μM unlabeled PFFs (green). No substantial contribution of R_ex_ is observed in either sample. The error bars in the relaxation data are propagated from the single exponential decay fitting errors of the underlying ^15^N-R_2_ data. All NMR data were acquired at 700 MHz ^1^H Larmor frequency.

### Off-pathway Ac-αS oligomers delay seeded fibril elongation

Aggregation of Ac-αS from free monomer results in a dynamic mixture of fibrils and off-pathway oligomers that are present simultaneously in solution (Figure 1a). We used the amyloid indicating ThT fluorescence change to investigate the interplay of PFOs co-existing with Ac-αS monomers and fibrils during seeded aggregation under conditions that promote elongation (10 mM PBS, pH 7.4). A significant delay in fibril formation is observed upon addition of 2x molar excess of big PFOs to PFF seeds. By adding big PFOs at a 5x molar excess to PFF seeds, amyloid formation is essentially abolished, as evidenced by the almost complete loss of ThT fluorescence intensity (Figure 6a). A similar inhibitory effect is also observed for the small oligomers (Figure S8c), indicating that fibril formation is significantly delayed in the presence of off-pathway oligomers. After five days of co-incubation with big or small PFOs, fibrils show morphology that is indistinguishable from the initial PFF seeds, which suggests that the PFOs modify and significantly delay the kinetics of fibril formation under conditions that favor templated elongation, but do not change the morphology of the mature fibrils produced (Figure S8d–f).

**Figure 6.**
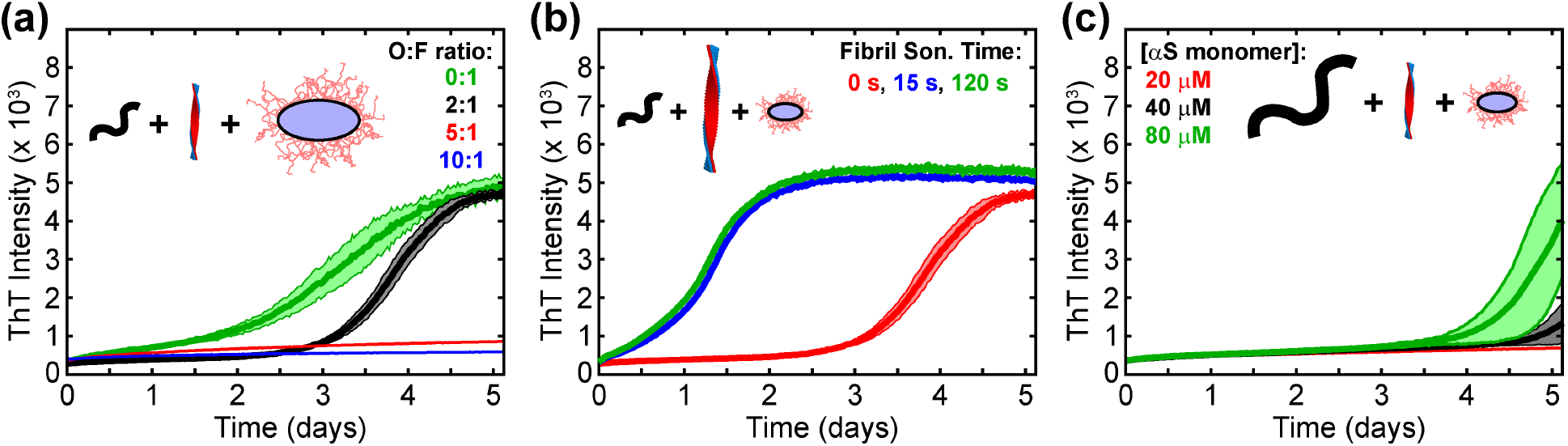
Off-pathway oligomers delay fibril seeding kinetics. Thioflavin T fluorescence assays of Ac-αS seeded aggregation under various conditions of the reaction components. (a) Seeded aggregation of 40 μM monomer, 1 μM (monomer equivalent) PFF, and either 0 μM (green), 2 μM (black), 5 μM (red), or 10 μM (blue) big PFO. (b) Seeded aggregation of 40 μM monomer, 1 μM PFF, and 2 μM big PFO, where the fibril seeds are sonicated for either 0 s (red), 15 s (blue), or 120 s (green) before addition to the reaction. (c) Seeded aggregation of either 20 μM (red), 40 μM (black), or 80 μM (green) monomer, with 1 μM PFF and 2 μM big PFO.

This inhibitory effect of the PFOs is reduced either by sonication of PFF seeds or by increasing concentration of monomers. Sonication of fibril seeds reduces the fibril length from micrometers to nanometers and produces more fibril ends, which provide more elongation sites for the same monomer-equivalent seed concentration (Figure S3a–b) (10, 46, 49). Consistent with the literature, the longer the sonication time applied to mature fibrils, and therefore the shorter the fibril seeds, the more numerous the elongation-competent ends and the more efficient the seeding capacity (Figure S3a–b, d–f). Using fibrils that have been sonicated for extended periods of time as seeds for 40 μM monomers, and co-incubating them with 2x molar excess PFOs relative to PFFs, the oligomer inhibitory effect on the aggregation kinetics is largely reduced (Figure 6b, S8a). Likewise, under the same conditions, increasing the monomer concentration corresponds to faster aggregation kinetics (Figure S3c, S8c). With increasing monomer concentration, we observe shorter lag times even in the presence of PFOs (Figure 6c, S3c, S8b). Thus under solution conditions favoring fibril elongation, promoting monomer–fibril interactions by increasing fibril ends or increasing monomer concentration reduces the inhibitory effect of off-pathway oligomers.

## 3. Discussion

### Identification of a monomer–fibril interface: N-terminal 11 residues of Ac-αS monomer interact with the fibril

Fibril seeding is an important process in the propagation and cell-to-cell spreading of disease related amyloidosis. Understanding the nature and residue specificity of monomer–fibril interactions is critical for understanding the mechanism of fibril seeding and for effective interference with aggregation processes. While numerous elegant experiments have focused on identifying regions in the monomer of Ac-αS that are important to the kinetics of fibril formation by studying point mutations and changes in primary amino-acid sequence or by altering solution conditions (28–30, 46, 50–52), here we present a molecular view of the binding interface between Ac-αS monomer and seeding-competent PFFs or seeding-incompetent PFOs. Our NMR relaxation based experiments reveal that the N-terminus of the free Ac-αS intrinsically disordered monomer forms an interaction interface with PFFs and PFOs under equilibrium conditions and under conditions in which there is excess monomer in solution. We demonstrate that PFFs and PFOs are dynamic entities that exist in equilibrium with monomeric Ac-αS (Figure 7). This is consistent with previous NMR translational diffusion results that indicated a population of monomer-like species in solution with Ac-αS fibrils (53) and with the concept of molecular recycling of amyloid fibrils first introduced with fibrils of the SH3 domain (54). Detailed ^15^N DEST experiments allow visualization of the transient binding interface of the monomer–PFF “dark-state” complex via the free, NMR observable monomer state. The NMR relaxation results indicate that the N-terminal 11 residues of the monomers bind transiently to the PFFs in equilibrium conditions and in fibril seeding conditions that promote elongation (Figure 7).

**Figure 7.**
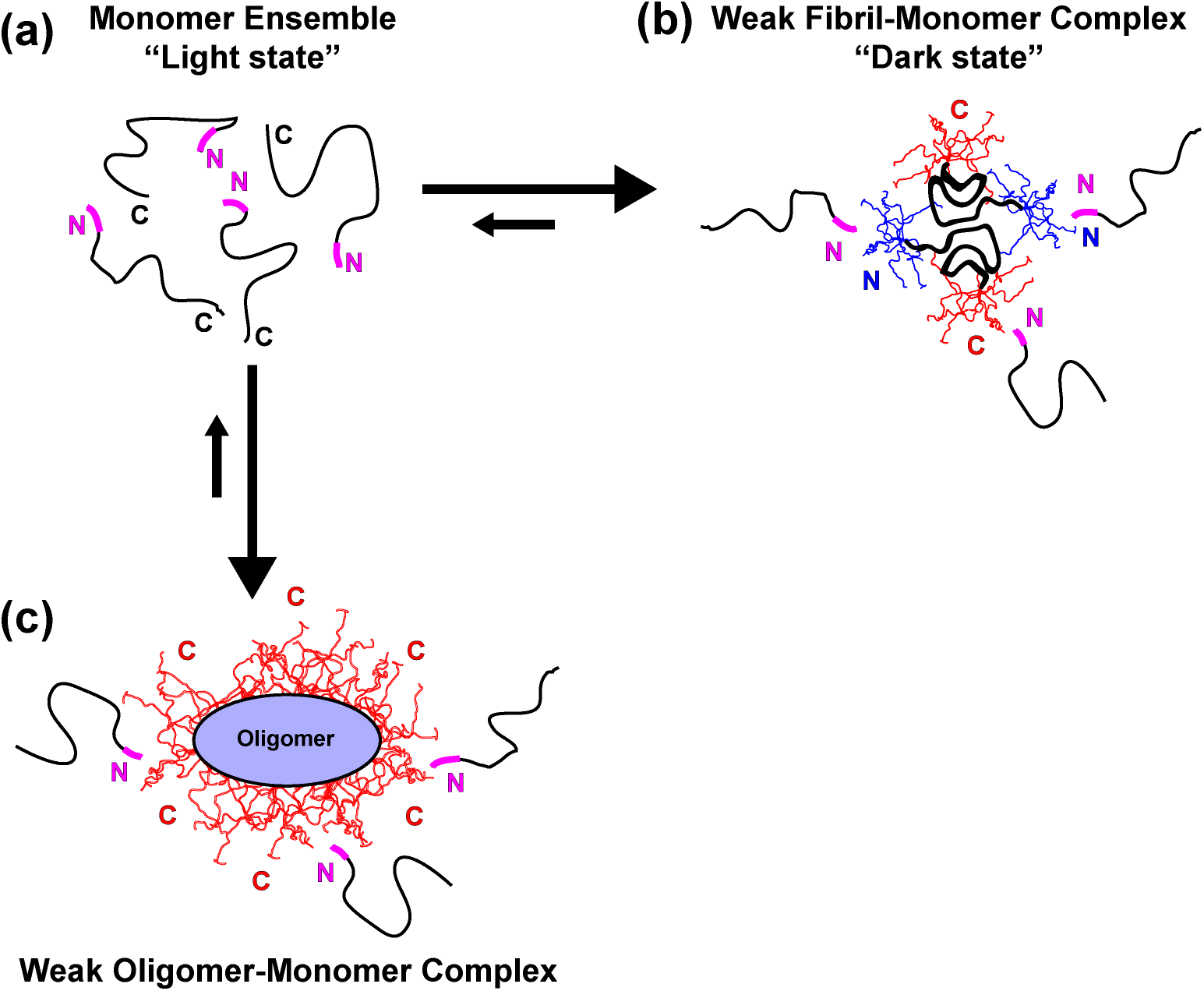
Proposed IDR–IDP interactions of the N-terminal monomer hot-spot with disordered flanking regions in the fibrils and oligomers. (a) Cartoon diagram of an intrinsically disordered unbound monomer ensemble. The flanking N-terminal residue interaction sites identified by ^15^N-DEST and ^15^N-R_2_ NMR experiments are represented in magenta. (b) Mature αS fibrils are in equilibrium with free monomer in solution. Based on the shared monomer binding site with off-pathway oligomers identified in this work and previous NMR studies on αS monomer–monomer interactions (66–69) we propose that Ac-αS monomers can interact via their N-terminal hotspot (magenta) with disordered flanking regions of mature fibrils. The fibril model is adapted from PDB: 6h6b (20), highlighting the disordered N-(blue) and C-termini (red) flanking the ordered Greek key amyloid core. This IDR–IDP interaction may drive monomer recruitment during amyloid seeding. (c) Off-pathway Ac-αS oligomers are also in equilibrium with free monomers, which re-associate with the oligomers via the same N-terminal hotspot (magenta). A schematic diagram of the an off-pathway oligomer based on the Otzen model (14) is shown, highlighting the disordered C-terminal cloud (red) surrounding the rigid core. We hypothesize that off-pathway oligomers delay amyloid formation by competing for interactions with the N-terminus of Ac-αS monomers.

### Importance of the Ac-αS N-terminus

Under physiological conditions *in-vivo*, the amino terminus of αS exists as the amino-terminal acetylated form. The N-terminal acetylation has been shown to have an impact on secondary structure propensity, metal binding, and aggregation kinetics (55–58). In addition, the N-terminus of αS has been recognized as an interaction hotspot that is important for membrane binding (59) and insertion (60) and may contribute to induction or inhibition of amyloid formation. N-terminal interactions of αS monomers with chaperones (61) and nanoparticles (62) have been shown to inhibit amyloid formation. Recently, a canonical motif for transient interactions of αS with multiple chaperones was determined (61), which shares the 11 N-terminal amino acids identified here. Together, this and previous work establish the N-terminus of αS monomers to be an important target for promoting or inhibiting amyloid growth.

### Ac-αS pre-formed oligomers are autoinhibitory: PFOs delay fibril formation in a seeding environment

We show that Ac-αS PFOs formed under fibril-promoting conditions are not merely bystanders in the aggregation process, but delay or even inhibit fibril growth under conditions that promote templated elongation. During the fibril formation process that favors elongation processes, many species including on-pathway and off-pathway oligomers, fibril seeds and mature fibrils exist in equilibrium. Our data, in which we characterize off-pathway PFOs, highlight that they can act as auto-inhibitors of Ac-αS fibril formation. Since the PFOs and PFFs share a common N-terminal interaction site on the monomer, we propose that the suppression of amyloid kinetics is due to a competition between the monomer N-terminus and the PFOs or PFFs (Figure 7). Competitive binding of monomers with PFOs results in reduced availability of free monomers to interact with PFFs, effectively inhibiting amyloid fibril elongation. Competitive mechanisms of suppressing amyloid formation have been previously proposed, such as competitive binding to fibril ends of Hsp70 to prevent fibril elongation (63) and high affinity monomer binders that sequester monomer to inhibit their participation in fibril growth (26). Inhibition of fibril growth by co-existing species has been proposed for other amyloid-forming proteins, such as insulin, lysozyme, and Aβ (64, 65). Our data shows how interactions between the multiple co-existing species even of the same protein may influence amyloid formation.

### Fibril seeding: proposed mechanism of monomer recruitment to fibrils

Based on our identification of a common N-terminal hotspot on Ac-αS monomers for interaction with both PFOs and PFFs, and on previous work by our lab (66, 67) and others (68, 69) on αS monomer–monomer interactions (Figure 7a,b), we propose a model in which the amino terminal residues of Ac-αS transiently interact with PFFs via the disordered flanking regions that surround the Greek key fibril core. Previous work has shown that transient IDP–IDP dimers are mediated by N–N or N–C terminal interactions (66–69). Specifically, NMR paramagnetic relaxation enhancement experiments in our lab highlighted that the early N-terminal residue A11, part of the monomer binding interface identified in the current study, preferentially forms intermolecular interactions with C-terminal residues in dimer formation (66).

Flanking IDRs in amyloid fibrils of multiple proteins have been shown to regulate protein and membrane binding of functional amyloids, modulate aggregation kinetics, and may play important roles fibril growth and monomer recruitment (27, 70). The remarkable similarity in the monomer recognition site for PFFs and PFOs is at first surprising, given the vast difference in morphologies between the PFFs and PFOs. However, similarity between the fibril and oligomer structures lies in their intrinsically disordered termini. Previous studies of oligomers purified under amyloid-forming conditions were determined to have an ellipsoid core with a fuzzy outer shell composed of disordered C-termini, consistent with our solution NMR data (14, 16, 47, 71). This “fuzzy-coat” model leaves only the C-terminus of αS oligomer exposed for interactions with monomers (Figure 7c). Conversely, αS amyloid fibrils possess both exposed disordered N- and C-termini flanking a rigid Greek-key fibril core, as observed in structural models determined by solid-state NMR and cryo-electron microscopy (19–23). In this way, αS fibrils also have a “fuzzy-coat” made of disordered C-termini (~residues 97-140) in addition to a “fuzzy-coat” made of disordered N-termini (~residues 1-36), providing the potential for αS monomers to interact with either the N- or C-termini of the fibrils (Figure 7b).

We therefore propose that the monomer–aggregate transient interactions that we have identified in this work occur between the amino terminal residues of the intrinsically disordered monomer and the intrinsically disordered flanking regions of the aggregates. While a substantial part of amyloid structures consist of intrinsically disordered regions in addition to the ordered core, the role and importance of these disordered flanking regions in amyloid seeding is not yet well understood, but is a current subject of interest (27). Here we have presented results that demonstrate that the monomer binding interface of the “dark-state” monomer–PFF complex resides in the amino terminal 11 residues of the monomer and lead us to hypothesize that these transient monomer–PFF interactions arise via intermolecular contacts between the N-terminus and the flanking regions of the fibril. These weak interactions may serve as the first step in the recruitment of monomers to fibrils. Thus, future therapeutic approaches for amyloid inhibition may consider targeting the IDRs that coat the fibril surface in order to block monomer recruitment and re-association with the fibril.

## 4. Materials and Methods (Condensed)

Please see SI Appendix for a detailed version of all materials and methods.

### Protein expression and purification

N-terminally acetylated human αS was obtained through co-expression with the pNatB plasmid (Addgene #53613) in *E. coli* BL21(DE3) cells. Uniformly ^15^N-labeled Ac-αS for NMR was expressed in M9 minimal media with ^15^N-ammonium chloride as the only nitrogen source. Unlabeled and [U-^15^N]-Ac-αS was purified as previously described (55). Protein purity and molecular weight were confirmed by ESI-MS. Protein was stored as lyophilized powder at −20°C until usage.

### Preparation of stable off-pathway oligomers and fibrils

Briefly, off-pathway oligomers were prepared by incubating 10 mg/mL Ac-αS powder in 10 mM PBS buffer (pH 7.4) at 37°C for 5 hours with shaking at 300 rpm. Stable oligomers were separated on a Superose 6 size exclusion column (GE Healthcare) at 4°C.

Fibrils were prepared as previously described (72). In short, fibrils were prepared by incubating 100 μL of 70 μM Ac-αS with a single Teflon bead (3 mm, Saint-Gobain N.A.) at 37°C, shaking at 600 rpm in 96-well clear bottom plates (Corning) in a POLARstar Omega fluorimeter (BMG Labtech) for at least 72 hours. To obtain varying lengths of fibril seeds, probe sonication was applied with 30% power.

### Experimental protocols

Detailed methods on thioflavin T fluorescence, transmission electron microscopy, circular dichroism, NMR, and cell toxicity experiments are provided in the SI Appendix.

## Data Availability

All data and protocols are available in the main text or the SI Appendix. Further inquiries can be addressed to the corresponding author.

## Acknowledgements

We thank Dr. M. Maral Mouradian (Rutgers Robert Wood Johnson Medical School) for access to cell culture equipment and materials and Mouradian lab members for experimental cell culture assistance. This work was funded by NIH grant GM110577 and GM136431 to J.B.

