## Supplemental Information for "NMR unveils an N-terminal interaction interface on acetylated-α-synuclein monomers for recruitment to fibrils"

**This PDF file includes:**

Supplementary Methods
Figures S1 to S8
SI References

### Supplementary Methods

#### Preparation of stable off-pathway oligomers and fibrils

Off-pathway oligomers were prepared by dissolving 10 mg Ac-αS lyophilized powder in 1 mL 10 mM PBS buffer (pH 7.4) and incubating the protein solution at 37°C for 5 hours with shaking at 300 rpm. The sample was then loaded on a Superose 6 size exclusion column (GE Healthcare) and eluted with PBS buffer at a flow rate of 0.3 ml/min at 4°C. Oligomer samples were concentrated with 3 kDa centrifugal filter units (Millipore Sigma) and were kept at 4°C until use. All oligomer samples were used within 24 hours. Fibrils were prepared as previously described (1). In short, protein powder was dissolved in 10 mM PBS (pH 7.4), and large aggregates were removed through centrifugal filtration using a 50 kDa molecular weight cutoff (Millipore Sigma). Fibrils were prepared by incubating 100 µL of 70 µM Ac-αS with a single Teflon bead (3 mm, Saint-Gobain N.A.) at 37°C, shaking at 600 rpm in 96-well clear bottom plates (Corning) in a POLARstar Omega fluorimeter (BMG Labtech) for at least 72 hours. Fibrils were washed multiple times with 10 mM PBS (pH 7.4) by re-suspension and centrifugation at 20k rpm for 30 mins in order to remove residual soluble and non-fibrillar components. Oligomer and fibril concentrations were measured by UV 280 nm after denaturation with 4 M guanidine hydrochloride.

#### Thioflavin T seeding experiments

Ac-αS protein powder was dissolved in 10 mM PBS (pH 7.4) and filtered through 50 kDa centrifugal units to remove large aggregates formed during lyophilization. Different concentrations of pre-formed fibril (PFF) seeds or pre-formed oligomers (PFOs) were added to 70 µM monomer solutions with 20 µM ThT. To obtain varying lengths of fibril seeds, probe sonication was applied to PFFs with 30% power. Morphologies and length distributions of sonicated PFFs were assessed by TEM. For each condition, 100 µL solution was dispensed into clear bottom 96-well plates and sealed with Axygen sealing tape (Corning). The plates were incubated at 37°C in a POLARstar Omega fluorimeter (BMG Labtech) under quiescent conditions. Fluorescence was monitored every 33 mins for at least 5 days.

#### Transmission Electron Microscopy (TEM)

3.5 µL samples were applied to carbon coated copper grids (Electron Microscopy Sciences) and incubated at room temperature for 90 s. Excessive solution on the grid was removed by gently tapping with filter paper. 3.5 µL 3% uranyl acetate solution (Fisher Scientific) was then added to the grid and incubated at room temperature for 60 s. The grid was then washed twice with 3.5 µL ultrapurified water. Images were recorded with a JEOL 1200EX electron microscope with 80k voltage.

#### Circular Dichroism (CD)

Far-UV wavelength spectra of PFOs and PFFs were measured on an AVIV 420SF CD spectrophotometer (AVIV Biomedical Inc.) in a 1 mm pathlength quartz cuvette at 25°C. Samples were diluted in 10 mM PBS (pH 7.4). Wavelength scans were obtained from 200–260 nm with a step size of 1 nm and 10 s averaging time.

#### NMR Experiments

All NMR experiments were performed in 10 mM PBS (pH 7.4) with 10% $D_2O$ at 4°C. To prepare [U-$^{15}$N]-Ac-αS monomer samples, lyophilized protein powder was dissolved in PBS buffer (pH 7.4), and large aggregates were removed using a 50 kDa centrifugal filter. [U-$^{15}$N]-Ac-αS PFFs were sonicated on ice by probe sonication at 30% power level with four cycles of 30 s power on and 30 s power off. Fibril morphologies were checked by TEM before and after NMR experiments. All concentrations reported are in monomer equivalents as determined by a BCA assay. All NMR data were processed using NMRPipe (2) and analyzed by Sparky (3).

Relaxation experiments. For all samples, $^{15}N$ transverse relaxation ($R_2$) experiments were measured from a series of HSQC-based 2D $^{1}H$–$^{15}N$ spectra using the Carr-Purcell-Meiboom-Gill (CPMG) pulse sequence with varying relaxation delays: [U-$^{15}N$]-Ac-αS PFFs and [U-$^{15}N$]-Ac-αS monomers at 800 MHz- 0, 8, 16, 16, 32, 64, 128, 192, 256, and 320 ms; [U-$^{15}N$]-Ac-αS PFOs and $^{15}N$-Ac-αS monomers at 800 MHz- 8, 16, 32, 48, 64, 96, 128, 160, 192, 224, and 256 ms; [U-$^{15}N$]-Ac-αS monomers alone or in the presence of PFFs at 700 MHz- 0, 8, 16, 32, 32, 64, 128, 192, 256, 320 ms; and [U-$^{15}N$]-Ac-αS monomers alone or in the presence of PFOs at 700 MHz- 8, 16, 32, 48, 64, 96, 128, 160, 192, 224, and 256 ms. The data were acquired by interleaving relaxation delays and $t_1$ increments. Experiments were acquired with a 1 ms interpulse delay ($\nu_{effective}$= 500 Hz). A single CPMG pulse train was constructed with eight 180° $^{15}N$ pulses. Protons were decoupled using a 180° $^{1}H$ pulse in the middle of each CPMG pulse train. The progressive solvent saturation effect was mitigated by introducing a $^{1}H$ steady-state pulse in the beginning of the sequence as implemented in ref. (4). A recycle delay of 2 s was used between each scan. $^{15}N$-$R_2$ rates were measured by fitting a single exponential decay function to peak intensities of the decay curves for each residue. Where reported, $\Delta R_2$ is the difference between the measured $^{15}N$-$R_2$ of the indicated sample and the $^{15}N$-$R_2$ of pure [U-$^{15}N$]-Ac-αS monomer at the same concentration. In the $^{15}N$-$R_2$ Hahn echo experiments (5) on [U-$^{15}N$]-Ac-αS PFFs and $^{15}N$-Ac-αS monomers at 800 MHz, the following relaxation delays were used: 0.768, 3.84, 7.68, 7.68, 19.2, 38.4, 76.8, 115, 154, and 230 ms. For $^{15}N$-$R_2$ Hahn echo experiments on [U-$^{15}N$]-Ac-αS monomers in the absence or presence of PFFs at 700 MHz, the following relaxation delays were used: 0.768, 7.68, 15.36, 30.72, 30.72, 46.08, 76.8, 107.52, 168.96, and 238.08 ms. The conformational exchange contribution to $R_2$ ($R_{ex}$) was determined for each residue as: $R_{ex} = R_2^{HE} - R_2$, where $R_2^{HE}$ is the apparent relaxation rate derived from the $^{15}N$-$R_2$ Hahn echo experiment.

*$^{15}N$-DEST experiments.* $^{15}N$-DEST experiments were conducted by applying an $^{15}N$ RF saturation pulse of 350 Hz or 150 Hz for 900 ms at variable $^{15}N$-frequency offsets: 350 μM [U-$^{15}N$]-Ac-αS monomer at 800 MHz- 0, ±0.5, ±1, ±2, ±4, ±8, ±18, and ±30 kHz; 350 μM [U-$^{15}N$]-Ac-αS PFFs at 800 MHz- 0, ±0.5, ±1, ±2, ±4, -8, +18, and ±30 kHz.. For each sample, a reference experiment in which 0 Hz RF saturation was applied at an $^{15}N$-offset of +30 kHz was also conducted. $^{15}N$-DEST profiles were obtained for each residue as peak intensity vs. $^{15}N$-offset. The ΔΘ profile was derived by calculating a Θ value for each residue, where $\Theta = \frac{(I_{30kHz}+I_{-30kHz})-(I_{1kHz}+I_{-1kHz})}{(I_{30kHz}+I_{-30kHz})}$, and $\Delta\Theta = \Theta_{(+)fibrils} - \Theta_{(-)fibrils}$. The residue-specific $\Delta R_2$ and DEST profiles of the [U-$^{15}N$]-Ac-αS PFFs were fit simultaneously to the McConnell equations using a simple two-state model in the DESTfit program (6, 7) to extract $p_{visible}$, $k_{on}^{app}$, and $R_2^{bound}$. The DEST profiles of 350 μM [U-$^{15}N$]-Ac-αS monomer were fit to a model in which the NMR-detectable state was not in exchange with the invisible state ($k_{on}^{app}$ = 0 $s^{-1}$) as a control.

**Cell culture**

Human SH-SY5Y neuroblastoma cells (ATCC) were cultured in DMEM/F12 (GE Healthcare) with 10% fetal bovine serum (FBS, Gibco Co.) and kept in a 5% $CO_2$ humidified atmosphere at 37°C. Cells were seeded onto 96-well (Corning) or 12-well glass bottom plates (Cellvis) for cell viability or immunocytochemistry experiments, respectively. Cells were allowed to recover for at least 24 hours, at which point they reached 75% confluency, before conducting experiments.

Cell viability MTS reduction assay. Cells were treated with different concentrations of PFFs or PFOs for 24 hours. Following the manufacturer's instructions, 20 μL 3-(4, 5-dimethylthiazol-2-

yl)-5-(3-carboxymethoxyphenol)-2-(4-sulfophenyl)-2H-tetrazolium reaction solution (MTS, Promega) was added to 100 µL cell culture and incubated for 2 hours at 37°C. Cell viability was measured as absorbance at 490 nm, which is directly proportional to the number of living cells in the culture.

_Immunocytochemistry_. Ac-αS samples (monomers, PFOs, and PFFs) were labeled with fluorescent ATTO488-NHS-ester (ATTO-TEC GmbH) per the manufacturer's procedure. In brief, samples were incubated with a 2 M excess of ATTO488-NHS-ester in labeling buffer (PBS/sodium bicarbonate solution, pH 8.3) for 1 h at room temperature. Unreacted fluorophore was washed from the conjugated ATTO488-PFFs by centrifugation at 20k rpm for 30 min and resuspension of the ATTO488-PFF pellet in fresh 10 mM PBS (pH 7.4); this washing procedure was repeated twice. Unbound fluorophore was washed from monomers and PFOs three times with 10 mM PBS (pH 7.4) by filtration with 3 kDa centrifugal units. Cells were treated with 3 µM ATTO488-Ac-αS monomers, PFOs, or PFFs and incubated for 24 hours.

Cells were fixed with 10% formalin (Sigma Aldrich) and permeabilized in PBS buffer with 0.5% Triton. Before being incubated with antibodies, cells were blocked with 5% Donkey Serum solution (Sigma Aldrich) for 30 min at 37°C. Cells were incubated with primary antibody (SYN-1, BD Biosciences) at 4°C overnight and protected from light. Afterwards, cells were washed with 10 mM PBS (pH 7.4) three times and incubated with fluorophore-conjugated secondary antibody (TRITC antibody, Sigma Aldrich) at room temperature for 1 h. Cell nuclei were stained with 4', 6-diamidino-2-phenylindole (DAPI, Millipore Sigma) for 90s and washed with PBS. All samples were imaged by a Zeiss LSM 780 confocal laser scanning microscope. Image analysis was done by Fiji (ImageJ) (8).

### Supplementary Figures

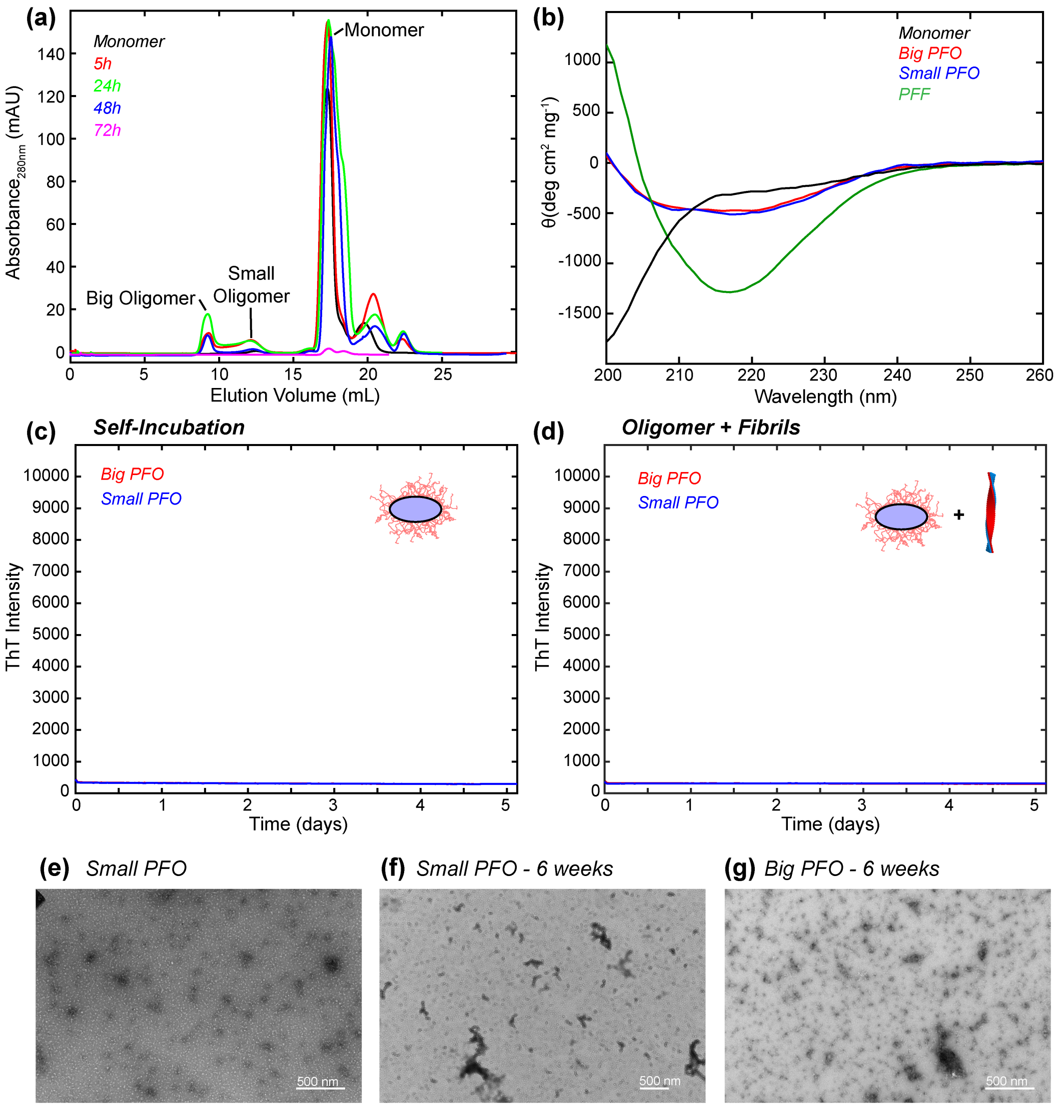


**Figure S1.** (a) SEC profile of 10 mg/mL Ac-αS monomer incubated at 37°C with shaking for 5 h, 24 h, 48 h and 72 h. Off-pathway oligomers do not convert into fibrils or larger aggregates with longer incubation times. After 72 h incubation, fibrils become the predominant species in the sample, greatly diminishing the oligomer and monomer peaks in the SEC profile. (b) Circular dichroism (CD) spectra of Ac-αS monomers, off-pathway pre-formed oligomers (PFOs), and pre-formed fibrils (PFFs) at 25°C in 10 mM PBS, pH 7.4 indicate their difference in secondary structure content. (c) ThT fluorescence assay of 40 μM big or small PFOs in 10 mM PBS, pH 7.4 incubated at 37°C in quiescent conditions. After 5 days, no increase in ThT fluorescence intensity was observed, indicating that these PFOs do not convert into fibrils. (d) ThT fluorescence assay of 40 μM big or small PFOs in the presence of 1 μM PFF seeds in 10 mM PBS, pH 7.4 incubated at 37°C in quiescent conditions. No increased ThT fluorescence signal was detected within 5 days, indicating that these off-pathway oligomers are not seeded by fibrils to propagate amyloid. (e) Representative TEM image of small PFOs. (f,g) Representative TEM images of small (f) and big (g) PFOs after 6 weeks of incubation at 37°C. Scale bars in (e)–(g) represent 500 nm. No amyloid fibrils are observed and oligomer morphology is maintained.

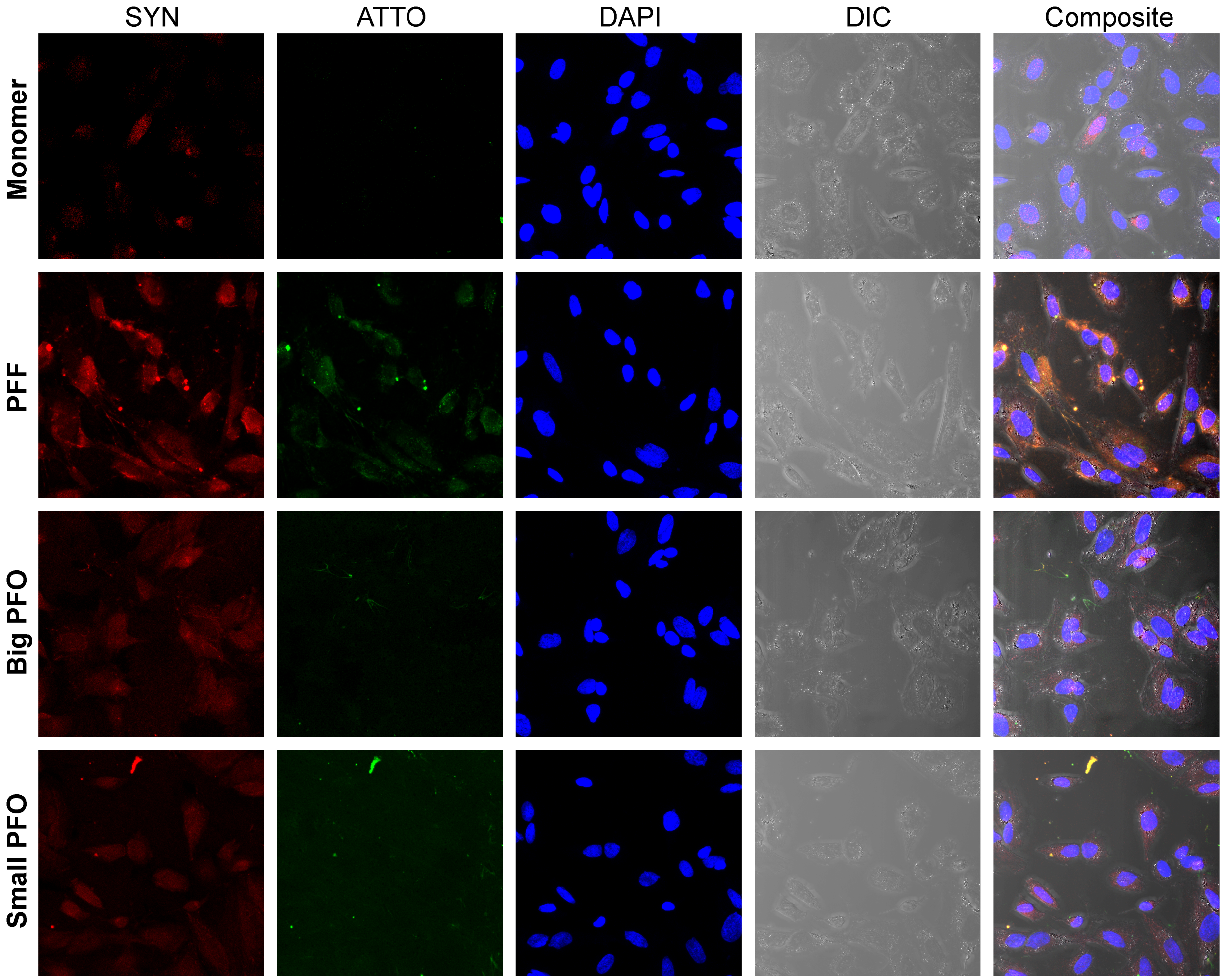


**Figure S2.** Confocal fluorescence images of SH-SY5Y cells treated with either monomer, PFF, big PFO, or small PFO, and then stained with anti-αS -antibody (red), ATTO dye (green), or DAPI (blue). The three separate channels corresponding to fluorescence signal from each dye are shown, along with the DIC image, which provides an outline of the cells. The final column is a composite image of all four of these channels.

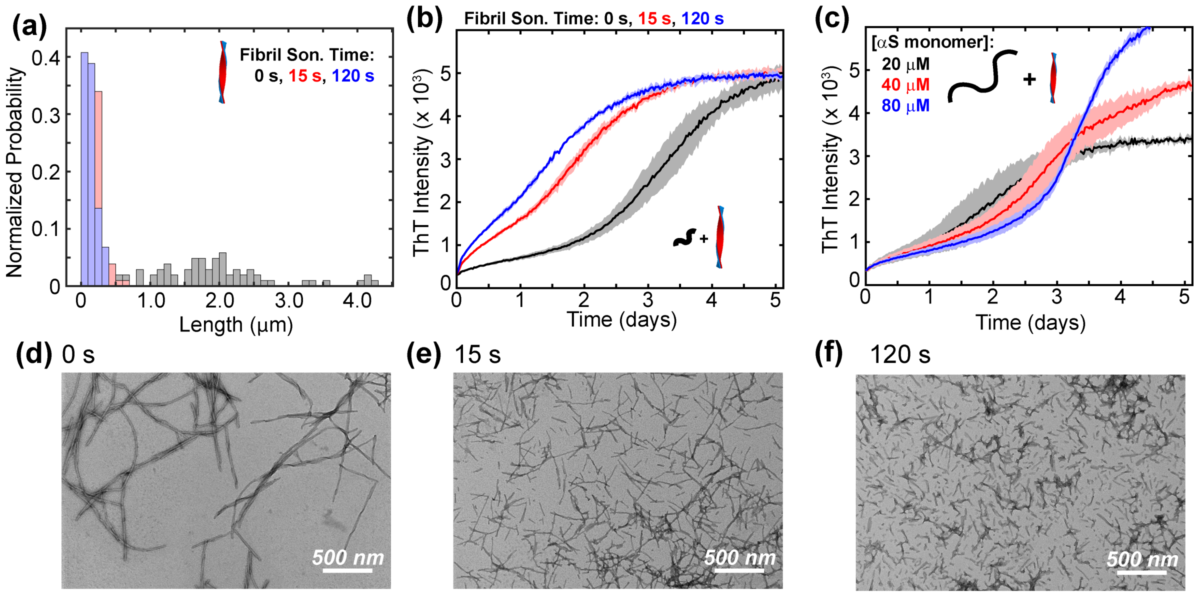


**Figure S3.** (a) The length distribution of Ac-αS fibrils as measured from TEM after sonication is applied at 30% power for different amounts of time as indicated. (b) ThT fluorescence assays of 40 μM Ac-αS monomer in the presence of 1μM PFF seeds that had been sonicated for varying amounts of time as indicated. (c) ThT fluorescence assays of varying concentrations of Ac-αS monomer in the presence of 1μM non-sonicated PFF seeds. All seeding experiments are conducted in 10 mM PBS, pH 7.4 at 37°C in quiescent conditions. (d–f) Representative TEM images of fibril seeds with (d) 0 s sonication, (e) 15 s sonication, and (f) 120 s sonication used for ThT assays.

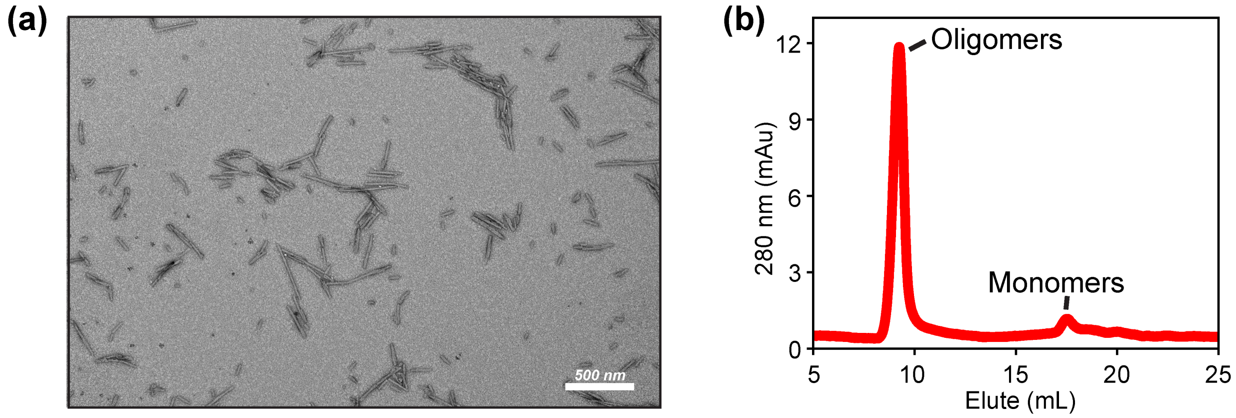


**Figure S4.** (a) Representative TEM image of sonicated fibrils used for NMR experiments in main text Figure 3a. Fibrils were prepared in 10 mM PBS, pH 7.4 at 37°C, shaking at 600 rpm as described in the Methods. Fibrils were then sonicated with a probe sonicator at 30% power for 120 s. (b) SEC profile of big off-pathway oligomers after re-injection. A low intensity monomer peak emerges, indicating that the oligomers are in equilibrium with monomers.

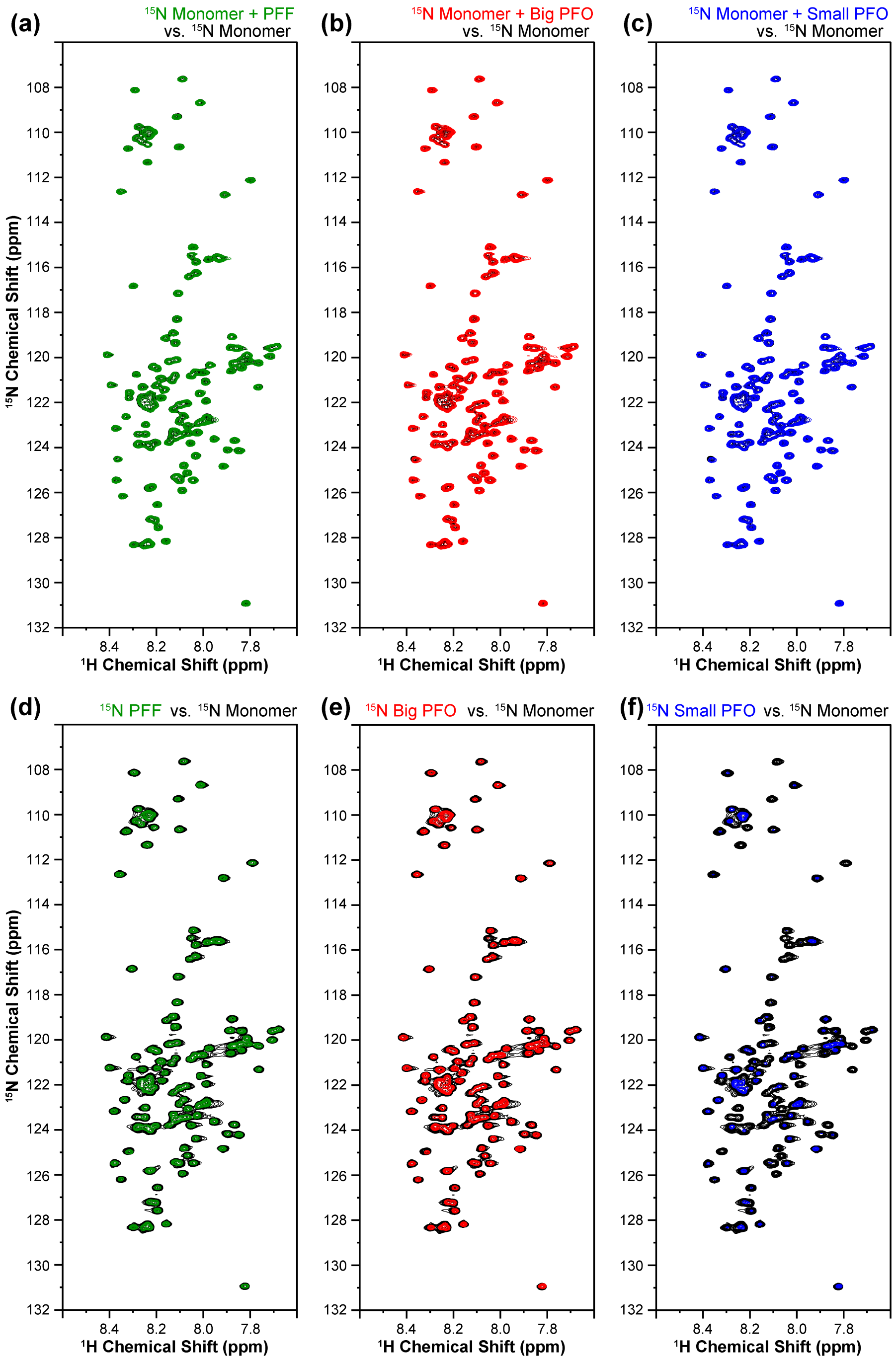


**Figure S5.** (a–c) $^{1}$H–$^{15}$N HSQC spectra acquired at 700 MHz $^{1}$H Larmor frequency of 90 μM [U-$^{15}$N]-Ac-αS monomer (black) overlaid with $^{1}$H–$^{15}$N HSQC spectra of 90 μM [U-$^{15}$N]-Ac-αS monomer in the presence of (a) 90 μM unlabeled PFF (green), (b) 90 μM unlabeled big PFO (red), and (c) 90 μM unlabeled small PFO (blue). (d–f) $^{1}$H–$^{15}$N HSQC spectra acquired at 800 MHz of 350 μM [U-$^{15}$N]-Ac-αS monomer (black) overlaid with $^{1}$H–$^{15}$N HSQC spectra of (d) 350 μM [U-$^{15}$N]-sonicated PFFs (green), (e) 350 μM [U-$^{15}$N]-big PFOs (red) and (f) 350 μM [U-$^{15}$N]-small PFOs (blue). No chemical shift perturbations are detected. All spectra were acquired in 10 mM PBS, pH 7.4 with 10% $D_2O$ at 4°C.

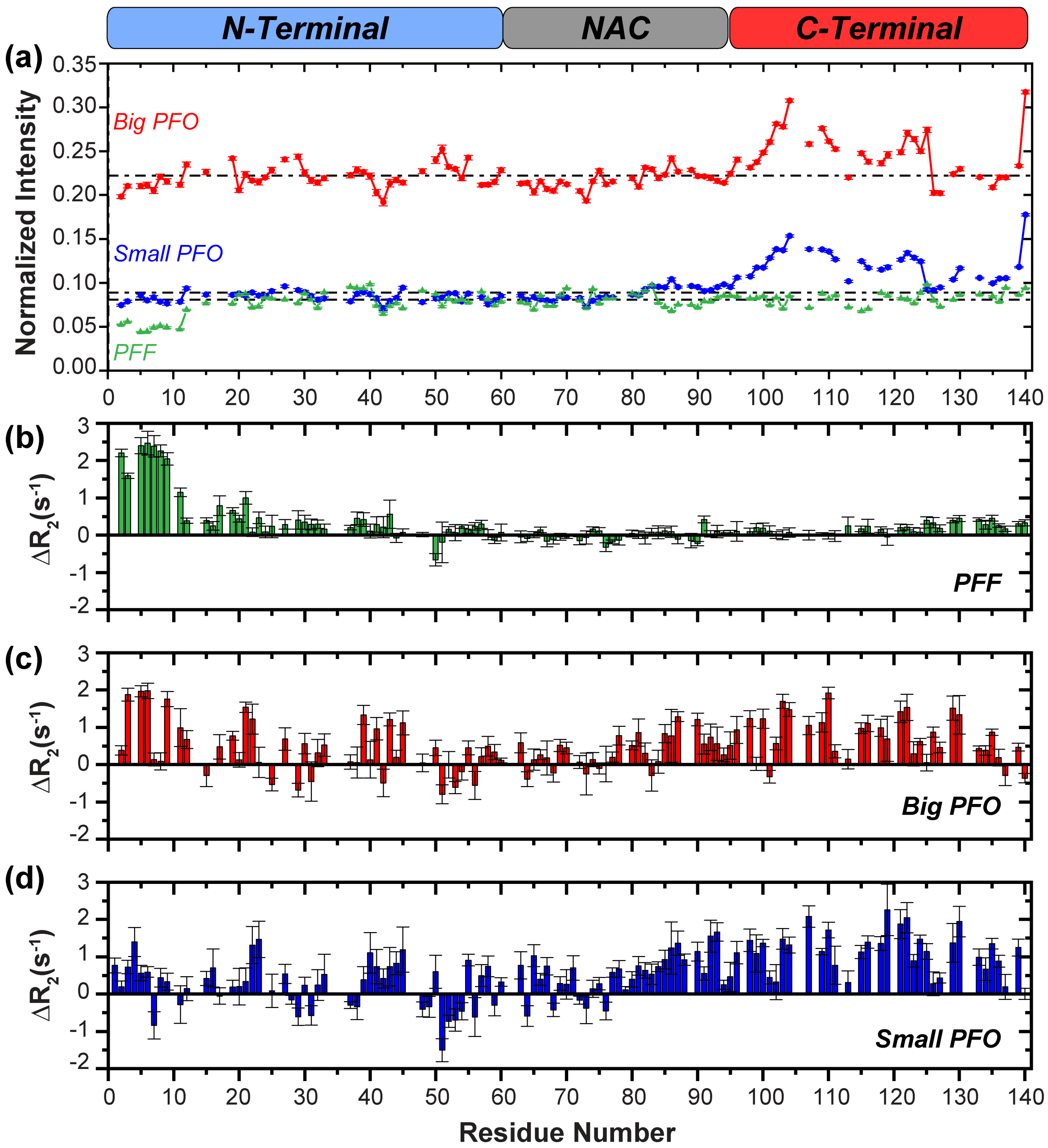


**Figure S6.** (a) Per-residue $^{1}H$–$^{15}N$ HSQC peak intensity ratios relative to 350 μM [U-$^{15}N$]-Ac-αS monomer of 350 μM [U-$^{15}N$]-big PFOs (red), 350 μM [U-$^{15}N$]-small PFOs (blue), or 350 μM [U-$^{15}N$]-sonicated PFFs (green). A dashed line is drawn at the median peak intensity ratio over the entire sequence for each aggregate species. The C-terminal residues of both PFO species show higher relative peak intensities than the median. Error bars are derived from the noise levels in each individual spectrum. (b–d) Residue-level $^{15}N$-$\Delta R_2$ rates of (b) 350 μM [U-$^{15}N$]-sonicated PFFs, (c) 350 μM [U-$^{15}N$]-big PFOs, or (d) 350 μM [U-$^{15}N$]-small PFOs calculated as $^{15}N$-$R_2^{aggregate}$–$^{15}N$-$R_2^{monomer}$. Error bars are propagated from single exponential decay fits of the underlying $^{15}N$-$R_2$ measurements. Together, higher peak intensities and $^{15}N$-$\Delta R_2$ rates in C-terminal residues of the PFOs are attributed in part to their disordered C-termini. All spectra were acquired in 10 mM PBS, pH 7.4 with 10% $D_2O$ at 4°C and 800 MHz $^{1}H$ Larmor frequency.

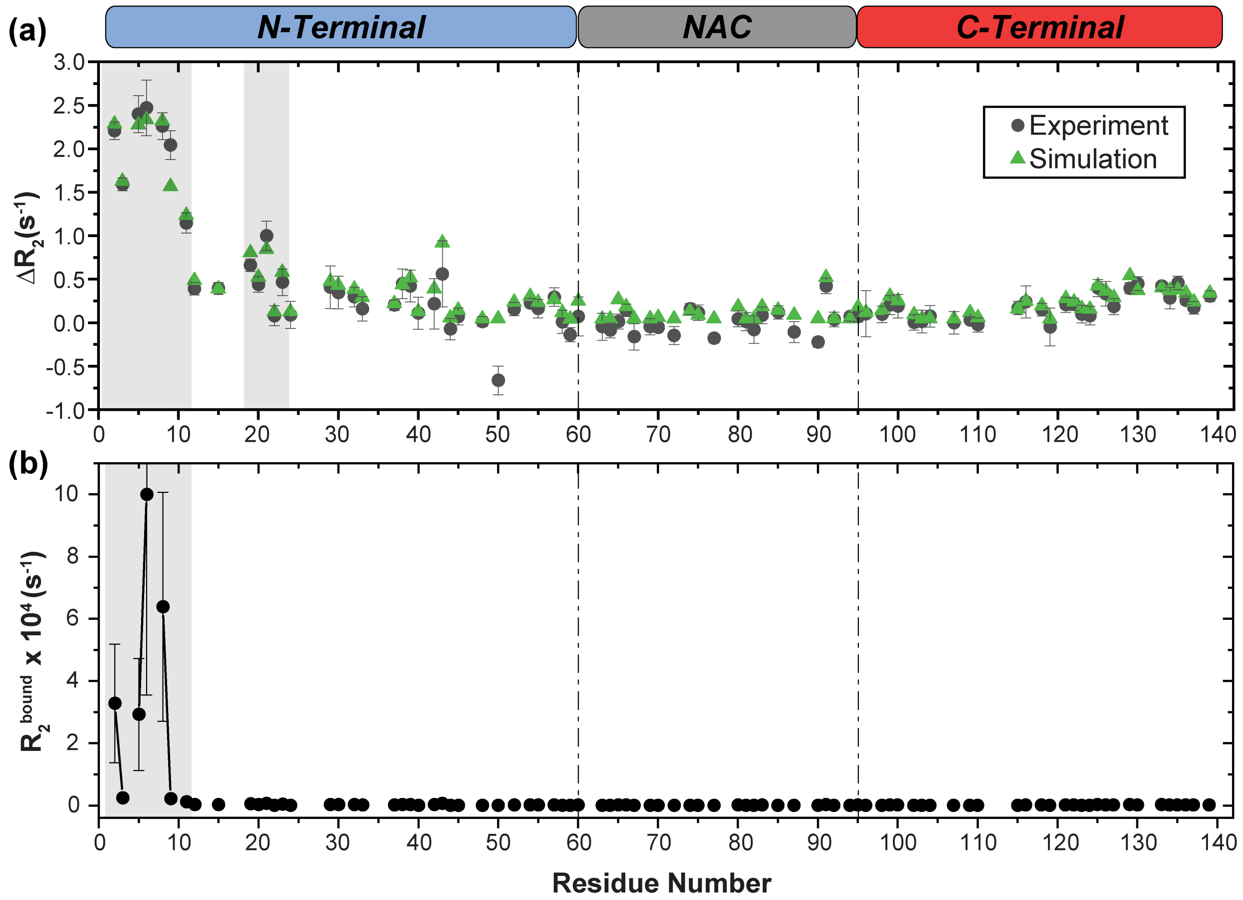


**Figure S7.** Simultaneous fitting of $\Delta R_2$ and $^{15}$N-DEST profiles of 350 μM [U-$^{15}$N]-Ac-αS sonicated PFFs to McConnell equations. (a) Black circles: per-residue experimental $\Delta R_2$ values calculated as $^{15}N\text{-}R_2^{fibril}$–$^{15}N\text{-}R_2^{monomer}$ (difference between $^{15}$N-$R_2$ data plotted in Figure 3b in the main text). Green triangles: Best fit $\Delta R_2$ values derived from simultaneous fitting of experimental $\Delta R_2$ and $^{15}$N-DEST profiles to a two-state model using the DESTfit program (6, 7). Regions in which consecutive residues exhibit $\Delta R_2$ values ≥ 0.5 $s^{-1}$ are highlighted in gray. (b) Per-residue $^{15}N\text{-}R_2^{bound}$ derived from simultaneous fitting of $\Delta R_2$ and $^{15}$N-DEST profiles to a two-state model using the DESTfit program (6, 7). Consecutive residues with an $^{15}N\text{-}R_2^{bound}$ ≥ 1000 $s^{-1}$ are highlighted in gray. Together, $\Delta R_2$ and $^{15}N\text{-}R_2^{bound}$ suggest that the first 11 residues of Ac-αS monomers directly interact with sonicated PFFs in equilibrium.

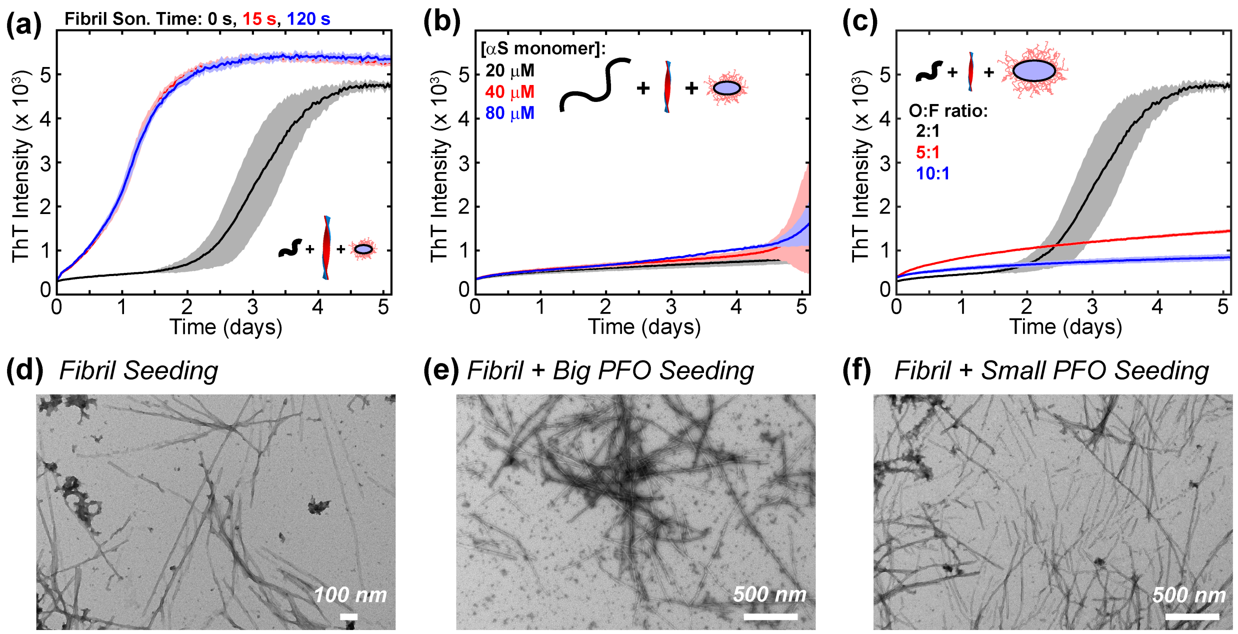


**Figure S8.** (a) ThT fluorescence assay of 40 μM Ac-αS monomers in the presence of 2 μM small PFOs and 1 μM PFF seeds that had been sonicated for varying amounts of time. (b) ThT fluorescence assay of varying concentrations of Ac-αS monomers in the presence of 2 μM small PFOs and 1 μM non-sonicated PFF seeds. (c) ThT fluorescence assay of 40 μM Ac-αS monomers in the presence of varying concentrations of small PFOs and 1 μM non-sonicated PFF seeds. All seeding experiments are conducted in 10 mM PBS, pH 7.4 at 37°C in quiescent conditions. (d–f) Representative TEM images of Ac-αS fibrils formed (d) in the absence of off-pathway oligomers, (e) in the presence of big PFOs, and (f) in the presence of small PFOs.
